# Profiles of circulating histidine-rich glycoprotein associate with chronological age and risk of all-cause mortality

**DOI:** 10.1101/464909

**Authors:** Mun-Gwan Hong, Tea Dodig-Crnković, Xu Chen, Kimi Drobin, Woojoo Lee, Yunzhang Wang, Fredrik Edfors, David Kotol, Cecilia Engel Thomas, Ronald Sjöberg, Jacob Odeberg, Anders Hamsten, Angela Silveira, Per Hall, Peter Nilsson, Yudi Pawitan, Sara Hägg, Mathias Uhlén, Nancy L. Pedersen, Patrik K. E. Magnusson, Jochen M. Schwenk

**Affiliations:** Science for Life Laboratory, Department of Protein Science, KTH - Royal Institute of Technology, Tomtebodavägen 23, SE-171 21 Solna, Sweden; Department of Medical Epidemiology and Biostatistics, Karolinska Institutet, Nobels väg 12A, SE-171 77 Stockholm, Sweden; Department of Public Health Science, Graduate School of Public Health, Seoul National University, 1 Gwanak-ro, Gwanak-gu, Seoul, 151-742, Korea; Department of Medicine Solna, Karolinska Institutet and Karolinska University Hospital, SE-171 76 Solna, Sweden; Cardiovascular Medicine Unit, Department of Medicine Solna, Karolinska Institutet, Akademiska Stråket 1, SE-171 64 Solna, Sweden; Department of Oncology, Södersjukhuset, Sjukhusbacken 10, SE-118 83 Stockholm, Sweden

## Abstract

Despite recognizing aging as risk factor of human diseases, little is still known about the molecular traits of biological age and mortality risk. To identify age-associated proteins circulating human blood, we screened 156 subjects aged 50-92 years using an exploratory and multiplexed affinity proteomics approach. We corroborated the top age-associated protein profile (adjusted P < 0.001) in eight additional study sets (N = 4,044 individuals), and confirmed a consistent age-associated increase (P = 6.61 × 10^-6^) by meta-analysis. Applying antibody validation determined circulating histidine-rich glycoprotein (HRG) as the target, and we observed that sequence variants influenced the antibodies ability to bind to the protein. Profiles of circulating HRG were associated to several clinical traits and predicted the risk of mortality during a follow-up period of 8.5 years (IQR = 7.7-9.3 years) after blood sampling (HR = 1.25 per SD; 95% CI = 1.12-1.39; P = 7.41 × 10^-5^). In conclusion, our affinity proteomics analysis found associations between the molecular traits of circulating HRG with age and all-cause mortality. This suggests that the profiles of multi-purpose protein HRG could serve as an accessible indicator of physiological processes related to aging.

## 2. INTRODUCTION

Aging is the single most dominant risk factor of common diseases in the elderly and of death in the human population (López-Otín, Blasco, Partridge, Serrano, & Kroemer, 2013). Molecular insights into aging could enable direct identification of future treatments for various diseases and would increase our understanding of longevity and related mechanisms. However, many of the underlying molecular processes and changes in humans remain poorly understood (López-Otín et al., 2013). Biological age or mortality risk have already been investigated via DNA methylation, telomere length, proteomic studies, mining of clinical records (Ganna & Ingelsson, 2015; Jylhävä, Pedersen, & Hägg, 2017) and showed several candidates for these traits (Barron, Lara, White, & Mathers, 2015; Ganna & Ingelsson, 2015; Marioni et al., 2015; Wiklund et al., 2010).

There are currently two major technological concepts available for measuring the proteins circulating in blood-derived samples: affinity-based proteomics and mass spectrometry. Both approaches offer a unique window into human health and diseases and have been used to study subsets of nearly 5000 protein circulating in blood (Schwenk et al., 2017). Affinity proteomics has initially suffered from a lack of binding reagents to the wider proteome, but antibody resources such as the Human Protein Atlas (HPA) (Uhlén et al., 2015) or aptamer-based platforms have enabled affinity proteomics for larger discovery projects, such as recently also in the context of aging (Lehallier et al., 2019). An important aspect for affinity proteomics is to validate the antibodies in a context dependent manner (Uhlen et al., 2016) and utilizing the power of population-based genome-wide association studies (GWAS) with circulating proteins (Suhre et al., 2017) can mitigated some of the uncertainty concerning target binding.

Utilizing antibody assays based on the suspension bead arrays (Byström et al., 2014), we profiled serum and plasma from a large number of individuals from different study sets. Studying the changes in age-related plasma protein levels, we explored, filtered and ranked plasma profiles associated with age across these sets of samples and confirmed antibody selectivity by genetic association tests and applying different immunoassays (Supporting Figure 1).

## 3. RESULTS

We profiled the serum proteomes of 156 humans to screen for age-associated proteins that could serve as indicators of biological age. The most significant finding was further investigated in additional 4,000 samples from eight different study sets (Table 1) and an approach utilizing different experimental methods and genomic data was used to validate antibody binding. The circulating protein profiles were examined in terms of different clinical traits and as a predictor of mortality risk which can reflect processed related to the biological age.

**Table 1.** Description of sample sets

| Study set | Age [yr] | Sex (F : M) | Sample Type | Indication* | Cohort name | References |
| --- | --- | --- | --- | --- | --- | --- |
| Set 1 | 50–92 | 78 : 78 | Serum | Population | TwinGene | (Lichtenstein et al., 2002; Magnusson et al., 2013) |
| Set 2 | 3–6 | 6 : 6 | Serum | Population | LifeGene | (Almqvist et al., 2011) |
|  | 9–63 | 102 : 102 | Plasma | Population |  |  |
| Set 3 | 48–93 | 1613 : 1386 | Serum | Population | TwinGene | (Lichtenstein et al., 2002; Magnusson et al., 2013) |
| Set 4 | 51–86 | 50 : 0 |  | Breast cancer |  |  |
| Set 5 | 56–75 | 0 : 50 |  | Prostate cancer |  |  |
| Set 6 | 55–78 | 16 : 27 | Plasma | Cardiovascular disease | IMPROVE | (Baldassarre et al., 2010) |
| Set 7 | 41–60 | 12 : 31 | Plasma | Myocardial infarction | SCARF | (Samnegård et al., 2005) |
| Set 8 | 48–73 | 20 : 23 | Plasma | Acute coronary heart syndrome | CHAPS | (Odeberg et al., 2014) |
| Set 9 | 40–73 | 600 : 0 | Plasma | Mammography | KARMA | (Gabrielson et al., 2017) |

\* Subjects included in the presented study did not include subjects diagnosed with the disease of the indication area, but the subjects assigned as controls for the different disease cohorts.

### Screening for age-associated profiles

Age-associated protein profiles were first investigated in a set of 156 human subjects selected in age intervals of 5 years from a Swedish twin cohort (denoted set 1) and as summarized in Table 1. The sex-matched samples included 30 monozygotic (MZ) twin pairs, who were 50 to 70 years old. Assays using a total of 7,258 HPA antibodies were applied to profile age-associated proteins in serum. The average intraclass correlation (ICC) within twin-pairs of antibody profiles was poor (ICC = 0.26), so twins were treated as unrelated. Minimal effects of the twin relationship were corroborated by a linear mixed model (LMM) that considered the dependency.

For this screening, target inclusion criteria were purely dependent on availability of antibodies, and not due to their target proteins. This set of antibodies comprised targets from 6,370 protein-encoding genes (about 32% of the non-redundant human proteome) and profiles were obtained using antibody suspension bead array assays (SBA). The acquired data was preprocessed and quality controlled, which included outlier removal and normalization to account for experimental variation across assay plates and data batches (details in Experimental procedures). Linear regression models (LM) then determined the protein profiles that changed monotonically with increasing age. The models revealed one out of 7,258 protein profiles age-associated when screening the sera of individuals at the ages of 50 to 92 years (adjusted P = 4.69 × 10^-5^). The association was also significant in the model considering twin-pairs (adj. P = 8.62 × 10^-5^).

### Replication of the discovered age-associations

Next, we continued to study the top finding in eight additional sample sets (set 2-9, Table 1). Details about the additional cohorts and sample selection are provided in the Supporting Information (see Study design and sample selection). We found consistent age-associated trends with HPA045005 across all eight replication sets (Supporting Figure 3). The combined effect of age on HPA045005 in all 9 sets was estimated using a random effects model accounting for differences in age ranges and distribution, showing a significant association between HPA045005 profile and age (meta-analysis, P = 6.61 × 10^-6^, Figure 1).

**Figure 1.**
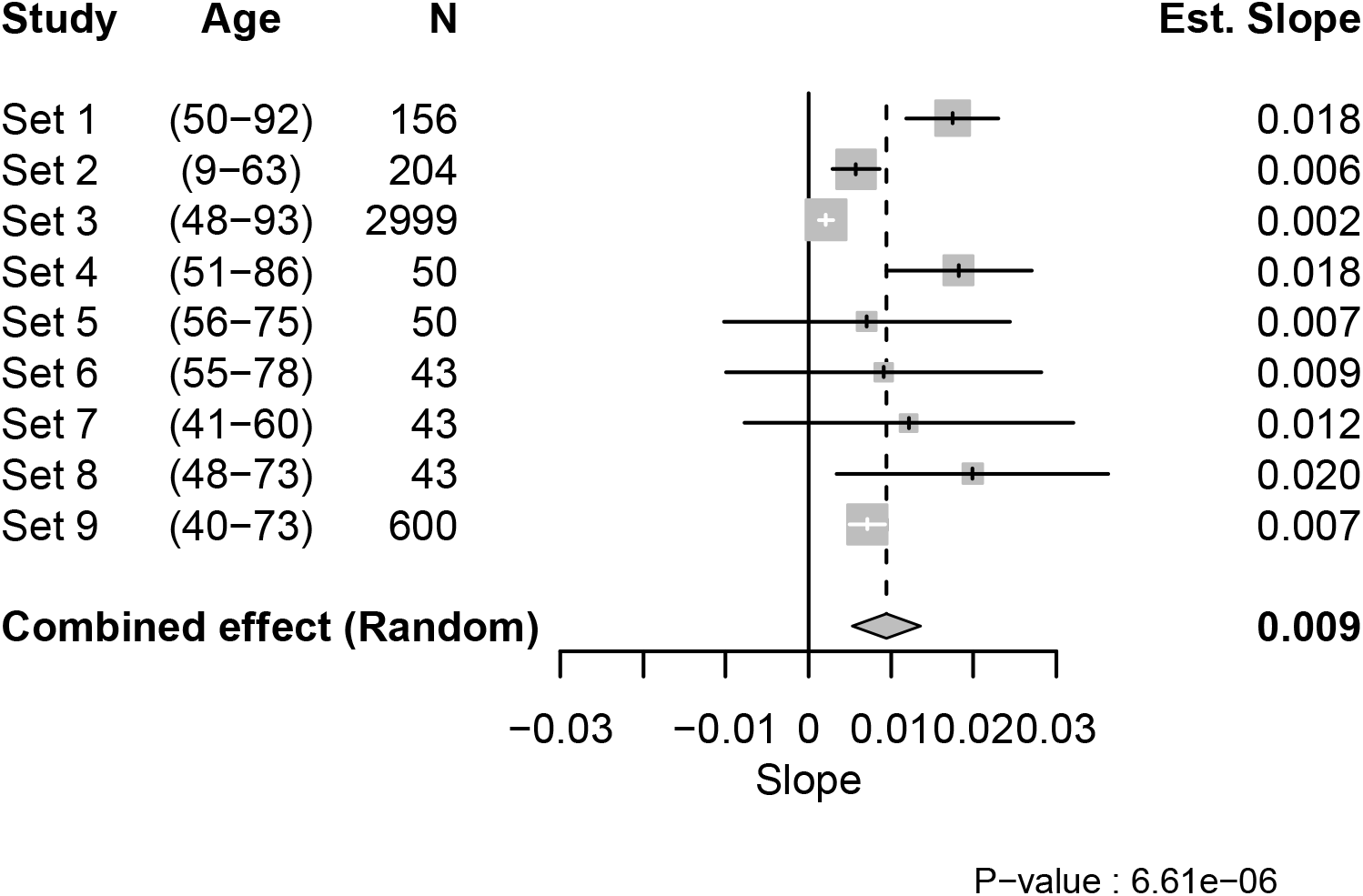
Meta-analysis from 9 different sample sets. In the forest plot, the numbers in parenthesis indicate the age range of the included subjects. For each sample set, the estimated effect from the linear regression model, 95% CI of it, and study weight in the meta-analysis are shown as a tick, a line, and a grey box, respectively, in the middle. The numeric value of the effect is clarified at the right side.

We then investigated a connection between genetic data and protein profiles obtained by HPA045005. A possible association in *cis* could infer about the circulating proteins captured by the antibody in our assays. Employing GWAS to ~8.8M genetic variants imputed from >700K single nucleotide variants genotyped by Illumina BeadChip in sample set 3 (N = 2308), we identified a single locus in chromosome 3q27.3 to be associated with the antibody profile (P < 1.13 × 10^-9^ = 0.01 / 8,833,947) (Figure 2A). The locus spans two genes, *FETUB* and *HRG*, in the human genome (Figure 2B). The most significantly associated genetic variant in the locus was the single nucleotide polymorphism (SNP) of rs9898 (P = 2.35 × 10^-97^, minor allele frequency (MAF) = 0.32). This SNP leads to an amino-acid change in the sequence of histidine rich glycoprotein (HRG) from Pro204 to Ser204 (pro-form). Among the SNPs at the top 99 genome-wide significance (P < 1.13 × 10^-9^) having RefSNPs (rs) number, four SNPs (rs1042464, rs2228243, rs10770) including rs9898 are non-synonymous and two SNPs (rs3890864 and rs56376528) are located near (<2 kbp) to the transcription start site (Supporting Table 1). All are located in exons or upstream of *HRG*. This GWAS result indicated that the antibody detected HRG, which is an abundant blood protein secreted by the liver (Uhlén et al., 2019). Associations of plasma HRG levels to SNPs, including rs2228243, have also been found in previous plasma profiling studies (Suhre et al., 2017).

**Figure 2.**
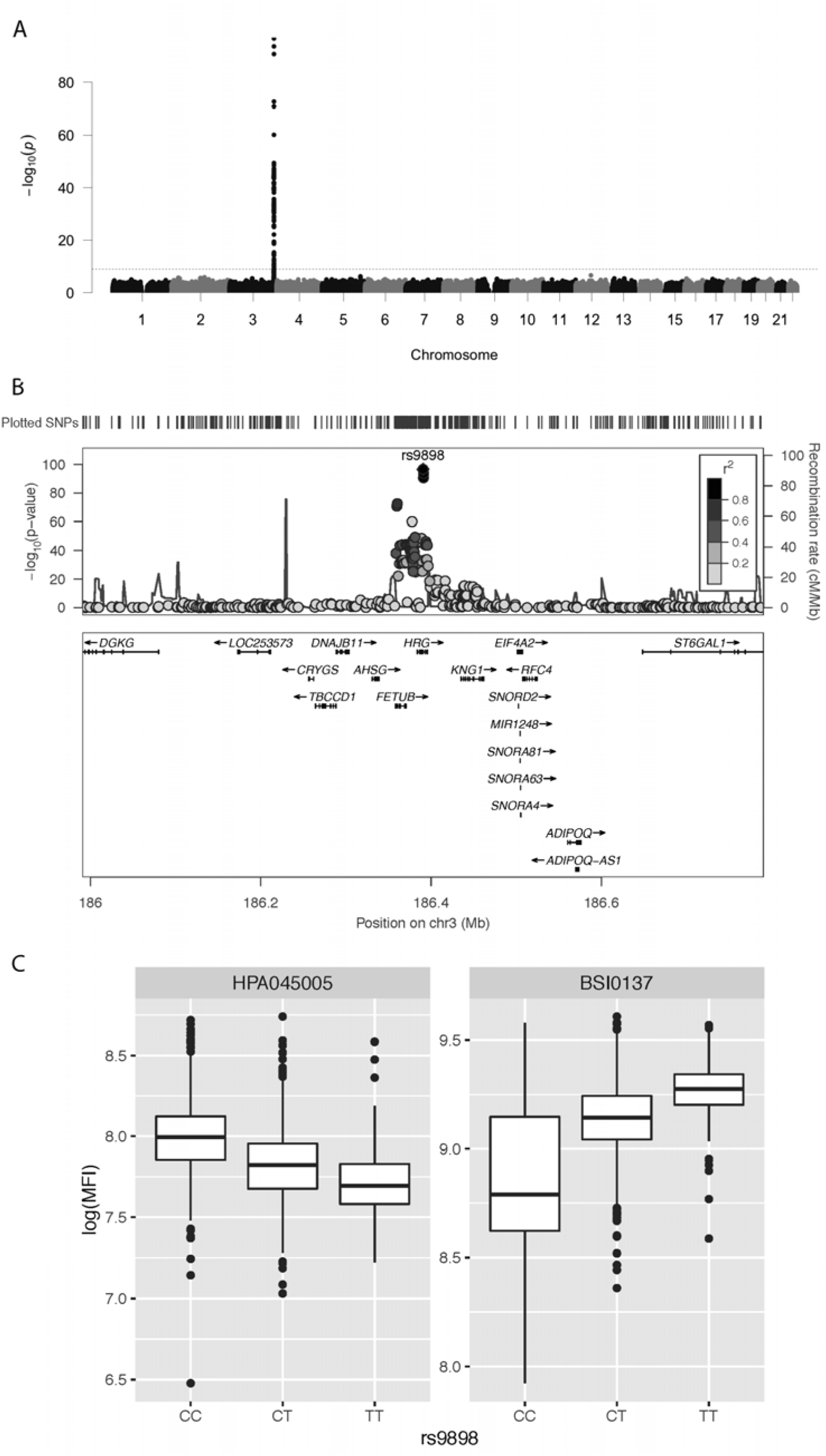
GWAS results of the age-associated plasma profile. (A) Manhattan plot. The significance of association between genotypes and HPA045005 profiles is presented vertically. The dashed guide line marks the stringent threshold of P-value for GWAS, which is P = 0.01 after Bonferroni correction. One peak in chromosome 3 indicates strong association of a locus with the molecular phenotype. (B) LocusZoom (Pruim et al., 2010) on associated locus. The illustration shows the elements of chromosome 3 associated with HPA045005 profiles. Zooming in on the peak of the Manhattan plot in (A), the genes around the locus are presented together with the associated SNPs. (C) Box plots to show the association between genotypes of rs9898 and two antibody profiles, HPA045005 and BSI0137. The trends were opposite.

### Validation and annotation of HRG profiles

Next, we confirmed the binding selectivity of HPA045005 to HRG and first used a bead-based sandwich immunoassay. Beads carrying HPA045005, another anti-HRG antibody (HPA054598), as well as negative controls were combined to detect full-length recombinant HRG in serial dilution. We found that pairing both HPA045005 and HPA054598 with a biotinylated version of HPA054598 allowed us to detect HRG in a concentration dependent manner (Supporting Figure 4A). Here, the curves obtained from both antibody pairs were substantially different and higher than the internal negative controls.

To elucidate the binding selectivity of HPA045005 also against other antigens, we applied the antibody to a large protein microarray. Among > 10,000 antigens, the antibody exclusively bound to its corresponding antigen (Supporting Figure 4B), which indicated that the antibody does not generally cross-react with other human antigens in an unspecific manner. We also aimed at determining the binding of HPA045005 to peptides representing its antigen using high density peptide arrays. The antigen did not reveal a significantly prominent recognition of these peptides above background (data not shown). Hence, binding analysis supports the findings from GWAS that HPA045005 captures HRG from serum and plasma in the single binder assay.

In addition to HPA045005, the GWAS with sample set 3 included several other antibodies of which one monoclonal binder (BSI0137) targeted the HRG protein. For BSI0137 there was one locus in the gene *HRG* which was strongly associated with the antibody’s profile (top pQTL was associated with P < 1 × 10^-300^, Supporting Figure 5). Interestingly, the identified locus included all the four non-synonymous SNPs previously observed to associate with HPA045005. However, the most significant SNP was not rs9898 but rs1042464, and the slopes of correlating BSI0137 with these SNPs were opposed to those for HPA045005 (Figure 2C). Applying Probabilistic Identification of Causal SNPs (PICS) (Farh et al., 2015), we confirmed that it was highly unlikely to observe rs9898 as the most significantly associated SNP with HPA045005 when rs1042464 was the causal SNP (none in 100,000 permutations), while the significance of rs1042464 was within possible range assuming rs9898 was causal (Supporting Figure 6). The distance from the mean of the permuted P values after log-transformation was about 1.22 standard deviation of the values. Likewise, the PICS applied for BSI0137 demonstrated that rs1042464 was much more likely to be causal than rs9898 for this antibody profile (Supporting Figure 6). The rs9898 causes HRG to contain either Pro204 or Ser204. Profiles of circulating HRG obtained from HPA045005 increased with the number of major allele C, which can produce only Pro204 form, in dosage dependent manner. Profiles of HRG reported by BSI0137 increased with the number of allele T of rs1042464 for Ile493. Hence, the PICS analysis revealed a selective binding affinity of HPA045005 towards another site of HRG than BSI0137. For these two binders, this points at differential selective affinities towards HRG variants: HPA045005 has a preference for HRG with Pro204 over Ser204 compared to BSI0137, which prefers HRG with Ile493 over Asn493. This suggested that the two antibodies preferred to recognized different isoforms of HRG related to partially correlating genetic variants.

Next, we investigated the associations of the HRG profiles to 10 clinical traits available in set 3. We applied a linear model that included the top SNP of each binder to account for the differences between the genotypes (Table 2). For HPA045005-derived HRG profiles, we observed negative associations to blood levels of hemoglobin (Hb), apolipoprotein A1 (APOA1), triglycerides (TG), and total cholesterol (TC), as well as a positive association to C-reactive protein (CRP). For BSI0137-derived profiles, there were significant associations to traits reflecting lipid metabolism such as levels of apolipoprotein B (APOB), low density lipoprotein (LDL), TC, TG, and expected negative associations to high density lipoprotein (HDL). This showed that different molecular traits could be associated to HRG depending on which binder and epitopes were used to report these. Additionally, we checked the association between the HRG profiles from HPA045005 and activated partial thromboplastin time (aPTT) in set 8. It was found negatively correlated with statistical significance (R = −0.55, P = 0.05).

**Table 2.**
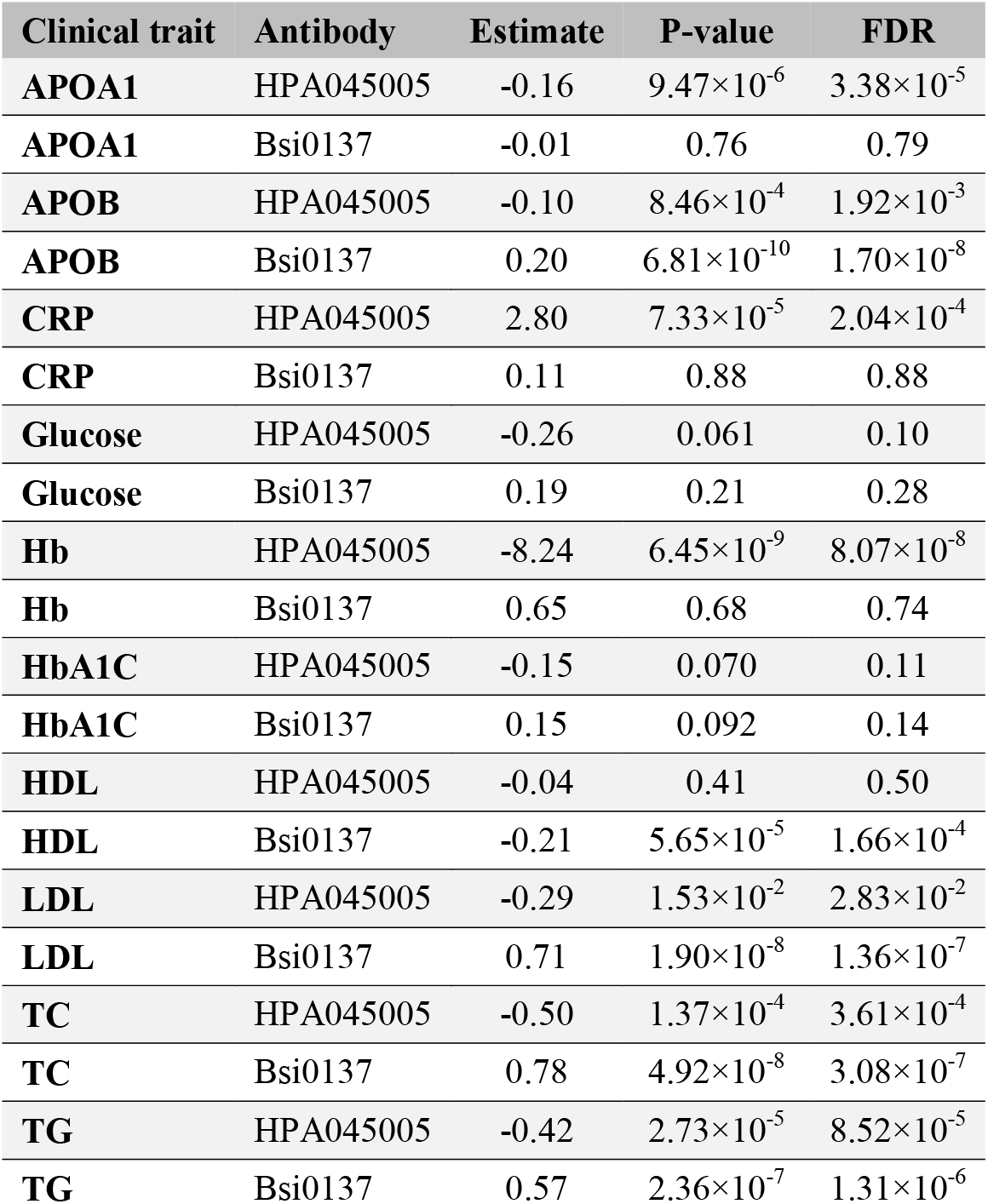
Associations of HRG profiles to clinical traits. Comparisons between association of HRG profiles to clinic traits using the linear models adjusted for age and the top SNP (rs9898 for HPA045005 and rs1042464 for BSI0137).

Lastly, mass spectrometry analysis of serum was carried out by LC-MS/MS to report peptides related to the HRG and to search for those representing variants rs9898, rs10770, rs2228243, and rs1042464. HRG peptides representing 96.5% of the sequence have been reported on PeptideAtlas (Desiere et al., 2006). HRG protein is an abundant blood protein of 30-100 μg/ml (Garcia et al., 1989; Schwenk et al., 2017), so no pre-fractionation or depletion of abundant proteins was performed prior analysis. One pool of serum was digested using five different proteases (Supporting Table 3) to increase the possibility that variant related peptides can be identified. As shown in Supporting Figure 8, the detected peptides represented the region of rs1042464 (Asn493 and Ile493). Additionally, data from the PeptideAtlas was used to search for evidence of the SNPs and peptide representing rs9898 were only found once as compared to >2800 times for rs1042464. However, peptides referring to rs10770 and rs2228243 were not detected and peptides related to rs9898 were very unlikely to be observed due to the altered amino acid composition around the cleavage site resulting in peptides of unsuitable peptide lengths.

### Mortality association and prediction

Finding the association of HRG and age led us to further study biological age in relation to mortality. We accessed the Swedish death registry for information on whether the subjects were still alive or not within a follow-up time of ~8.5 years (IQR = 7.7-9.3) after donating blood. We chose the largest sample set of the subjects at mid to old ages that spanned the average life expectancy in Sweden, which was sample set 3 (N = 2973, 48-93 years old). This was to gain statistical power needed for the analysis of all-cause mortality, which was otherwise a relatively rare event. A Cox proportional hazards model with age as the time scale was used with the adjustment for the effects of sex. This revealed that the profiles from HRG obtained by HPA045005 were significantly associated with the mortality risk during follow up (P = 1.13 × 10^-4^). The HRG profiles determined by BSI0137 were scarcely correlated with those of the HPA045005 (R^2^ = 0.006) and not associated to mortality (P = 0.57).

To adjust for the effect of chronological age at sampling on the mortality association of HRG profiles derived from HPA045005, the data was standardized using a linear model for age and age squared for each sex separately. The hazards model using the standardized HRG value affirmed the association with mortality (P = 7.41 × 10^-5^). The risk of all-cause mortality was estimated to increase 1.25 times per standard deviation (SD) of the age and sex adjusted HRG values (95% confidence interval (CI) = 1.12-1.39). In the model accounting for potential genetic effect of the most significantly associated SNP rs9898, the estimated HR of HRG became even higher with similar significant (HR = 1.31 per SD, P = 9.02 × 10^-5^, N = 2,307). No evident difference was observed when stratifying by the genotype of the SNP on HR of HRG (Supporting Table 2).

A Cox model stratified by sex suggested stronger mortality association in women (HR = 1.36 per SD, P = 2.13 × 10^-4^) than in men (HR = 1.15 per SD, P = 0.059). Comparing extreme subsets with standardized HRG levels of the upper and lower quartile demonstrated that the difference of median age at death was 1.8 years in favor of the bottom quarter (P = 4.60 × 10^-3^, HR = 1.53, Figure 3). The difference was 1.9 years in men (86.9 years vs. 85.0) and 0.6 years in women (89.6 vs. 89.0; Supporting Figure 7). The difference in life expectancy between the two extreme quarters at the age of 45 was 3.6 years in women (87.4 vs. 91.0) and 2.7 years in men (83.9 vs 86.6), assuming age-at-death follows a Weibull distribution (Supporting Table 4).

**Figure 3.**
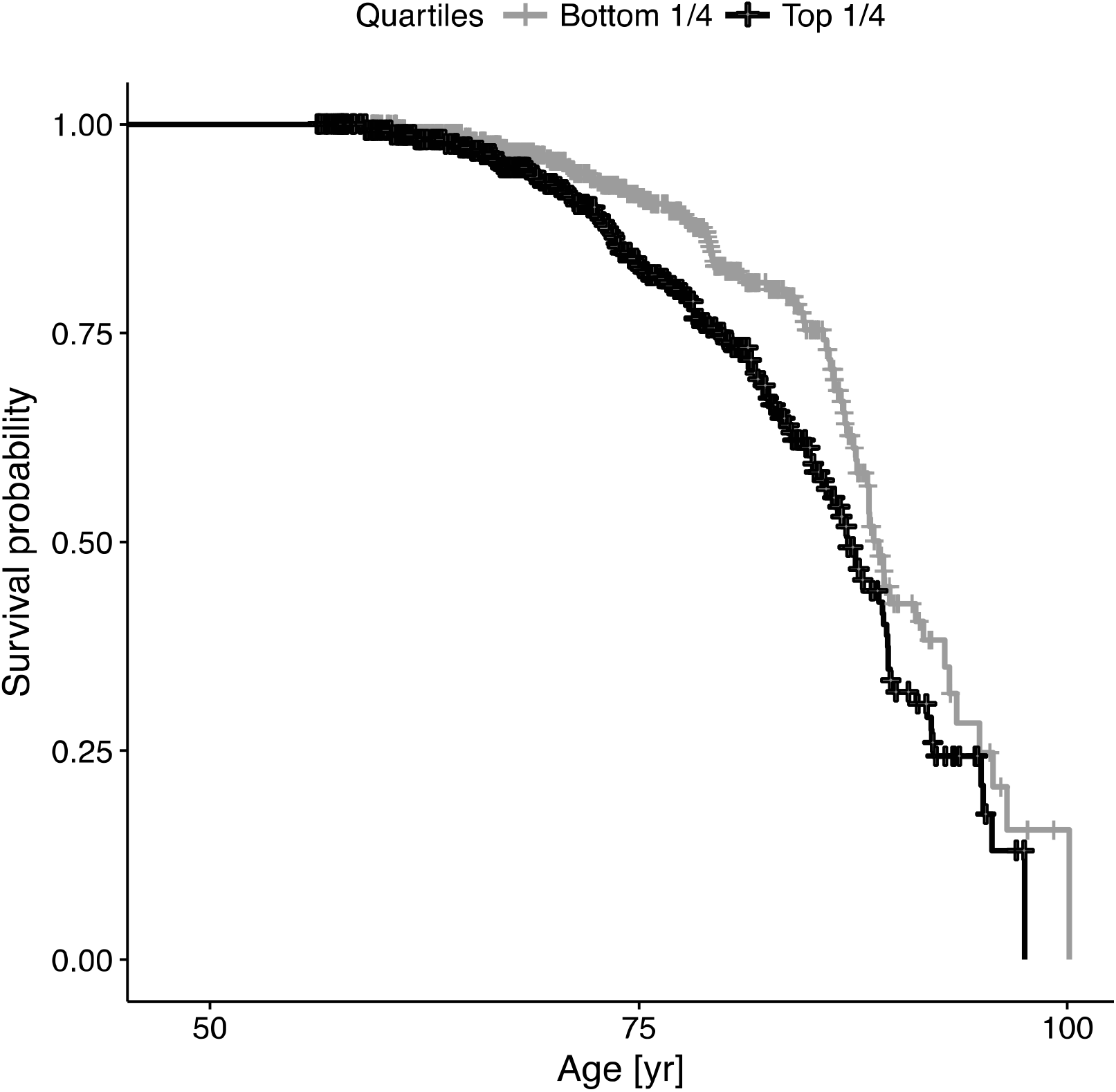
Survival analysis comparing upper and lower quarters of HRG levels. The individuals of sample set 3 were divided into four subsets by the quartiles of HRG levels. Differential mortality across follow-up time is illustrated by the survival curves. Some detailed statistics related to this survival analysis are presented in Supporting Table 3.

The potential influence of inflammation on survival was also tested including clinically measured CRP. As for HRG, two Cox models were fitted using 1) CRP and 2) age-adjusted CRP levels. The latter was obtained using the same linear model as HRG and adjusting for the same covariate. Associations of CRP model 1 (HR = 1.07 per SD, P = 0.023) and model 2 (HR = 1.01, P = 0.023) therefore less pronounced than for HRG. Next, we included CRP as a covariate in the Cox model for HRG to determine if inflammation in general would have an influence of HRG-related mortality. Negligible attenuation of the HRG association was observed after additional adjustment for CRP (HR = 1.25 to 1.24 per SD, P = 6.45 × 10^-5^ to P = 1.20 × 10^-4^). We also assessed the relation of HRG profiles to age and mortality considering diabetes related traits to account for any short-term effects of glucose. The results of the Cox models adjusted for 1) Glucose and 2) HbA1c were barely changed for both models (HR = 1.25 to 1.25 per SD for both, model 1: P = 6.45 × 10^-5^ to P = 4.40 × 10^-5^, model 2: P = 4.05 × 10^-5^, respectively). For all those survival analyses, we confirmed that none of the hazard models violated the proportionality assumption of the Cox model using Schoenfeld residuals (Grambsch & Therneau, 1994). A summary of the survival analyses with 95% CI is presented in Supporting Table 5.

## 4. DISCUSSION

### HRG is a multi-functional protein circulating in blood

We analyzed the age-related proteins in blood by multiplexed antibody-based assays and found a consistent, positive association between HRG profiles and age. Validating the binding of the antibody using GWAS, protein microarrays and sandwich assays revealed that HRG was captured from serum or plasma. We further demonstrated that a selective binding affinity towards variants of HRG revealed associations with mortality risk as compared to HRG variants preferred by another anti-HRG antibody.

HRG has been described as an abundant protein in human blood plasma and according to mRNA sequencing data of human tissues, HRG is exclusively expressed in liver (Uhlén et al., 2015; Uhlén et al., 2019). The protein has been characterized to interact with diverse molecules including heparin, heme, immunoglobulin G, Zn^2+^, and complement components (Poon, Patel, Davis, Parish, & Hulett, 2011; Priebatsch, Kvansakul, Poon, & Hulett, 2017) and particular functions have been assigned to each of its six domains (Martin et al., 2018). HRG is involved not only in immune response toward foreign substances and clearance of dead cells, but also in vascular biology including anti-coagulation (Poon et al., 2011). HRG levels have previously been correlated and linked to blood ABO type and age (Drasin & Sahud, 1996). The protein has been named a biomarker of preeclampsia, which entails angiogenic imbalance and defective coagulation control (Bolin, Akerud, Hansson, & Akerud, 2011). Partly due to its molecular composition and abundance, HRG has been assigned to many other different biological processes. When searching for possible protein-protein interactions of HRG using the STRING database (Szklarczyk et al., 2019), we observed that the number of listed interactions were enriched for platelet degranulation (Gene Ontology:0002576). Many of the proteins in the HRG-interactome were also expressed by the liver and secreted into blood (Uhlén et al., 2019).

With the current data, it remains difficult to postulate the most plausible mechanism why HRG increased in the process of aging and why this observation was only seen for one of the two anti-HRG antibodies. However, HRG has functional similarities with CRP, which is another indicator of aging and mortality (Barron et al., 2015). Their related functions in the context of coagulation and inflammation (Poon et al., 2011), may increase the likelihood that both change with age due to similar reasons. Interestingly, we found a negative association of the HRG profiles of HPA045005 to levels of blood hemoglobin, which could otherwise point at the involvement of this HRG in binding free heme from erythrocytes. Observing a negative correlation between the HRG profiles and aPTT, for which a prolonged time indicates lower thrombosis risk, supports a hematological hypothesis: lower HRG profile might indicate lower risk of thrombosis, and thereby contributes to reducing the risk of mortality. In contrast, the profiles of HRG affected by variants on residue 493 did reveal links to lipid metabolism but no significant association to Hb levels. This supported the hypothesis that a site- and variant-specific binding of the two antibodies (via distinct epitopes) revealed different perspectives about the traits of HRG.

In the *HRG* gene, there are 4 relatively common genetic variants (minor allele frequency > 10%) that lead to amino acid polymorphisms. In a genetic study investigating the aPTT, Houlihan et al. observed that the minor T allele of rs9898 was associated with shorter aPTT, suggesting an elevated risk of thrombosis in these individuals (Houlihan et al., 2010). Houlihan et al. proposed a potential interaction between HRG and thrombosis, hence thrombosis could be a possible mediator between elevated HRG and risk of mortality. The genetic association seems opposite to the two correlations we observed here, 1) more T allele and lower HRG profiles of HPA045005 and 2) lower HRG profiles and longer aPTT. However, considering the correlation of the HRG profiles with the rs9898 was mainly driven by molecular characteristic of our immunoassay, the opposite trend strengthened the idea of the association of the HRG profiles with mortality through not rs9898 but thrombosis. Tang et al confirmed Houlihan et al.’s observations and pointed out that possible interactions between causal variants of *HRG*, *KNG1* and *F12* may further influence coagulation and aPTT (Tang et al., 2012). We also performed a look-up about the variants using the data hosted by the Genotype-Tissue Expression (GTEx) Project portal (version 8, accessed 2019-11-19). We searched for any significant expression QTL (eQTL) or splice QTL (sQTL) that could provide further insights about circulating levels of the secreted HRG. For the rs9898 (as well as rs2228243 and rs10770) there were however neither any eQTLs nor sQTLs reported in GTEx. The rs9898 does not alter the mRNA expression or associated with the formation of splice-variants of *HRG*.

GWAS analysis revealed the associations of both antibodies’ profiles with those non-synonymous SNPs. The PICS analysis provided indications about which amino acid residue of the HRG may affect the antibody recognition. Using this novel approach, we found that HRG profiles from HPA045005 differentiated between HRG variants with Pro204 over Ser204. This might imply that the antibody had its HRG binding epitope in proximity to residue 204. This position is located in HRG’s cysteine protease inhibitor domain for which a lacking inhibitory activity was previously described (Ochieng & Chaudhuri, 2010). It is noteworthy that no single genetic variant around *HRG* reached genome-wide significance for mortality risk in a study including the TwinGene cohort (Ganna et al., 2013). Interestingly, two of the neighboring residues to Ser204 or Pro204 are amino acids with functional groups: Asn202 is a N-glycosylation site found for Ser204 variants (Hennis et al., 1995) and Cys203 has been identified as a site for glutathione modification (Kassaar et al., 2014). Investigating the functional differences of these isoforms may assist to further categorize HRG’s involvement in a diverse set of physiological processes.

### Limitation and variation

Circulating HRG was profiled across ages and samples from different donors to cover a broad range of lifespans. Even though finding consistent trends of HRG in serum and plasma of the cross-sectional study, the performance of the HRG assays can be influenced by the type of blood preparation. This could influence the degree to which HRG associated with age. On the other hand, gradual associations were repeatedly observed in multiple independent study sets derived from different Swedish cohorts, which provided supportive indications about the association of HRG with age. As profiles of circulating HRG increased as age advances, an analysis of longitudinal samples collected from representative subjects could be a viable strategy. As we also found that the elevated HRG was correlated with higher risk of mortality, the age-dependent diversity may imply a time-wise transition along individual ages, possibly biological ages. Lastly, molecular investigations are needed to further interpret the increasing trends of the HRG profiles, and as suggested by the differential associations to the clinical traits, to confirm the physiological differences of the various HRG isoforms.

We observed variation in the degree to which HRG associated with age, which is visible in Figure 1. To some extent, the variation can be explained by the shift of signal range in each study set, which was primarily developed to screen for possible associations and not standardized to determine absolute abundance levels. Seeing that the estimated slopes from sample sets 2, 3 and 9 were relatively lower than the values from the other sets, some parts of the variation might originate from the difference in age range and sample source, sample collection and preparation, or selection of participants. For example, the individuals in the sample sets 2, 7 and 9 were substantially younger (median age 40, 52 and 54 years, respectively) compared to all others (~65 years old). The sample sets 2, 3, and 9 were near population-based, while the others were healthy individuals except those in sample set 1, in which older women and men were overrepresented due to same number of individuals selected per age group.

### Significance of HRG as a potential predictor of mortality

Several other molecular indicators have been previously reported to predict mortality risk. Barron et al. showed that CRP (HR = 1.42), N-terminal pro brain natriuretic peptide (NT-proBNP, HR = 1.43), and white blood cell (WBC, HR = 1.36) count were statistically significant in meta-analyses (Barron et al., 2015). The HR estimate of HRG in our study (1.53 between top and bottom quarters) was comparable. Schnabel et al. linked CRP to mortality risk in a follow-up period of median 8.9 years (Schnabel et al., 2013). Using HR per SD, their estimate for CRP was 1.18, which was similar to our estimate of 1.25 from HRG. Marioni et al.’s HR per SD of DNA methylation (1.09 – 1.21) was also comparable to our HRG estimate (Marioni et al., 2015). Ganna and Ingelsson used questionnaire-derived measures for an extensive population-based mortality study (Ganna & Ingelsson, 2015). Their top predictors resulted in Harrell’s C-index = 0.74 when including age, and using the same model, HRG performed in the same range (C-index = 0.77).

Indeed, other proteins like GDF15, IL-6 and CRP have previously been described in aging and all-cause mortality independent of telomere length (Wiklund et al., 2010). We discuss GDF15 and its relation to medication in the Supporting Text, because it exemplifies that other, possibly unknown factors or unavailable data can assist to explain the cause of an increase in the observed HRG profiles. We acknowledge that a better understanding of HRG in the context of aging and mortality will require more lifestyle data from the donors. Lastly, our presented strategy was of an exploratory nature and we did not actively include antibodies towards previously known age-related proteins, such as GDF15. The shortlisted targets represented those for which the antibodies performed in the antibody bead array method when screening serum and plasma for indicators of aging.

In conclusion, we have described circulating HRG profiles as indicator of aging and mortality. Extensive efforts were put into confirming our observation across independent cohorts and applying molecular approaches to characterize the differential recognition of HRG. As a known multi-purpose adapter protein, HRG plays a role in hemostasis and the profiles of the protein can serve as a predictive indicator for all-cause mortality within 8.5 years after blood draw. Profiles of circulating HRG could be helpful as an accessible indicator of processes linked to biological aging.

## 5. EXPERIMENTAL PROCEDURES

### Cohort design and sample selection

#### a) Sample set 1 from TwinGene cohort

A population wide collection of blood from 12,614 twins born between 1911-1958 has been undertaken in a project called TwinGene. The primary aim of the TwinGene project has been to systematically transform the oldest cohorts of the Swedish Twin Registry (STR) into a molecular-genetic resource (Magnusson et al., 2013). From 2004 to 2008, a total of 21,500 twins (~200 twin pairs per month) were contacted by the invitation to the study containing information of it and its purpose, also consent forms and health questionnaire. The study population was limited to those participating in the Screening Across the Lifespan Twin Study (SALT) which was a telephone interview study conducted in 1998-2002 (Lichtenstein et al., 2002). Other inclusion criteria were that both twins in the pair had to be alive and living in Sweden. Subjects were excluded from the study who had declined to participate in future studies or been enrolled in other STR DNA sampling projects. When the signed consent forms returned, blood-sampling equipment was sent to the subjects, who were asked to visit local health-care facilities on the morning, after fasting from 20:00 the previous night, from Monday to Thursday and not the day prior to a national holiday. This was to ensure that the sample tube would be delivered to the Karolinska Institutet (KI) Biobank by the following morning by overnight mail. After arrival, the serum was stored in liquid nitrogen.

The contribution for sample set 1 of serum samples from the TwinGene study consisted of: A) samples from 96 unrelated twins distributed in groups of 12 subjects (6 males and 6 females) in each age strata 50, 55, 60, 65, 70, 75, 80 and 85 years of age, and B) samples from 60 MZ twins (30 complete pairs) distributed in groups of 12 (3 male pairs, 3 female pairs) in each age strata of 50, 55, 60, 65 and 70 years of age. The width of the age intervals was approximately ±3 months.

#### b) Sample set 2 from LifeGene cohort

LifeGene is a prospective cohort study that includes collection of plasma and serum, tests of physical performance, as well as questionnaire responses regarding a wide range of lifestyle factors, health behaviors and symptoms (Almqvist et al., 2011). Participants respond to a webbased questionnaire and book time for a visit to a LifeGene test center, at which blood samples are taken. EDTA plasma was processed at the test center as follows: the EDTA tube with a gel plug was centrifuged, put into −20°C prior to shipment in a cold chain. All samples were sent to KI Biobank for further separation into aliquots in REMP plates and frozen at −70°C. All participants or, in the case of children under the age of 11, their guardians, provided signed consent.

The sample set 2 cohort consisted of 5 male and 5 female samples randomly chosen from each of the ages <5, 10, 15, 20, 25, 30, 35, 40, 45, 50 and 55 (±3 months). For 12 participants, serum was also available.

#### c) Sample sets 3-5 from TwinGene cohort

Sample sets 3, 4, and 5 were selected from the same cohort, TwinGene (Magnusson et al., 2013), as for sample set 1 (described above). Out of 132 microtiter 96-well plates for storage of TwinGene samples, the twelve plates having the largest age span (>20 years) among samples in a plate and another randomly chosen twenty plates comprising a sufficient number of samples (>91) were selected. Sample set 3 consisted of the three thousand samples in the selected 32 storage places. The data of one individual was removed in the analyses because age of the subject is missing. Independently from the sample selection, sample sets 4 and 5 were age and gender matched controls for breast and prostate cancer studies, respectively. The mortality data was obtained by connecting individuals in TwinGene to the data in the Swedish tax authorities by personal identification number. The data was updated on 2015-01-10. Clinical blood chemistry assessments were performed by the Karolinska University Laboratory for the following biomarkers: total cholesterol, triglycerides, HDL, LDL (by Friedewald formula), CRP, glucose, APOA1, APOB, Hb, and HbA1c (Magnusson et al., 2013). Clinical blood chemistry assessments were performed by the Karolinska University Hospital Laboratory. Levels of HbA1c were measured by a high-liquid performance chromatography separation technique. Levels of the other biomarkers were determined by Synchron LX systems (Beckman Coulter) (Rahman et al., 2009).

#### d) Sample sets 6-9

The sample sets 6 to 9 are described in Supporting Text.

#### e) Ethics

All the studies were approved by the Ethics Board of the correspondent hospital or institution, and conducted in agreement with the Declaration of Helsinki. The ethical approval document numbers are 2007/644-31/2 for TwinGene, 2009/615-31/1 for LifeGene, 03-115 and 2017/404-32 for IMPROVE, 95-397 and 02-091 for SCARF, EPN 2009/762 and LU 298-91 for CHAPS, and 2010/958-31/1 for Karma. All subjects, or their guardians, provided their informed consent to participation in individual studies.

### Plasma proteomics analysis methods

Experimental details on the molecular methods and data processing are available as Supporting Information. This included descriptions for the antibody selection, SBA, sandwich immunoassays, mass spectrometry and protein microarray analysis

### Genome-wide association study

Genomic DNA from all available dizygotic twins and one member of each monozygotic twin pair were genotyped by using Illumina OmniExpress BeadChip (700K). Genotyping QC exclusion criteria: genotypic or individual missingness > 0.03, MAF < 0.01, Hardy-Weinberg equilibrium P < 10^-7^, sex mismatch, heterozygosity (individuals with an F-statistic beyond ±5 SD from the sample mean), or cryptic relatedness. The 1000 Genome reference panel (GRCh37/hg19, Phase 1, version 3) was used for imputation, by using Mach 1.0 and Minimac. After genotype antibody-profile match, GWAS was performed among 2308 twins by using PLINK 1.90 beta. Analyses were restricted to autosomal SNPs with imputation quality (info or r^2^) > 0.4. The first four principal components were included to control population stratification. The --within option in PLINK was used to statistically adjust for relatedness (complete dizygotic twin pairs). Manhattan plots were drawn using qqman package in R 3.4.1. The mutation types of the associated SNPs were obtained from UCSC table browser (https://genome.ucsc.edu) using human ‘GRCh38/hg38’ assembly and ‘snp150Common’ (dbSNP build 150, ≥1% MAF) table, which was accessed on 2018-09-20.

### Statistical analysis

The preprocessed intensity data was log-transformed ahead of downstream analyses. To control family-wise error rate, the Bonferroni method was employed for adjusting P-values unless otherwise specified. The ICC was computed using Shrout and Fleiss’s method for a set of randomly chosen two raters (Shrout & Fleiss, 1979). The linear association of an antibody signal level with age was tested with ordinary LM using R 3.6.0. The meta-analysis was conducted using the inverse variance method with between-study variance estimated by DerSimonian-Laird model (DerSimonian & Laird, 1986) with “meta 4.9.9” R-package. We used a LMM to address the correlation between twins where the response variable was the normalized antibody measurement and age was a fixed covariate. This model was performed using the R-package “lme4 1.1.21”. For the association test for mortality, Cox proportional hazards models were fitted to the survival data with age as the time-scale and right censoring of the age on the updated date of death information (Thiébaut & Bénichou, 2004). In the survival analysis for two group comparison, the subjects in sample set 3 were divided into two groups, top and bottom quarters by the standardized HRG values, which were the scaled residuals of LM where the normalized MFIs of HRG were regressed on age and age squared for women and for men, separately. The hazard models were adjusted for sex if applicable and for CRP, glucose, or HbA1c as described above. The proportionality assumption of the models was tested using Schoenfeld residuals (Grambsch & Therneau, 1994). Survival analyses including computation of Harrell’s C-index (Harrell, Califf, Pryor, Lee, & Rosati, 1982) were conducted using the R packages “survival 3.1.8” and “eha 2.8.0”.

## 6. ACKNOWLEDGMENTS

We like to thank Camilla Björk and Jens Mattsson from MEB at the Karolinska Institutet, everyone in the Affinity Proteomics group at SciLifeLab, and especially Claudia Fredolini, MariaJesus Iglesias, Matilda Dale, Sanna Byström, Martin Zwahlen, Björn Forsström, Björn Winckler, and Philippa Pettingill for supporting this work. We also thank the entire staff of the Human Protein Atlas for their efforts, Hanna Tegel and Johan Rockberg and their team for providing the recombinant HRG protein. We thank Bernd Wollscheid from ETH in Zurich for fruitful discussions about HRG. This work was supported by ProNova VINN Excellence Centre for Protein Technology (VINNOVA, Swedish Governmental Agency for Innovation Systems), the Knut and Alice Wallenberg Foundation for the Human Protein Atlas, Science for Life Laboratory for Plasma Profiling Facility, Erling Persson Foundation for KTH Centre for Precision Medicine. We acknowledge The Swedish Twin Registry for access to samples and data. The Swedish Twin Registry is managed by Karolinska Institutet and receives funding through the Swedish Research Council under the grant no 2017-00641. LifeGene was supported by grants from the Swedish Research Council, Torsten and Ragnar Söderbergs Foundation, Stockholm County Council, and AFA Försäkringar.

## 8. AUTHOR CONTRIBUTIONS

M-G.H. and J.M.S. designed the study. T.D-C., K.D., F.E., D.K. and R.S. acquired and analyzed the proteomic data. M-G.H., X.C., and Y.W. analyzed GWAS data. M-G.H., T.D-C., W.L., C.E.T, Y.P., S.H., P.K.E.M., and J.M.S. performed and discussed the statistical analyses. J.O., A.H., A.S., P.H., N.L.P., and P.K.E.M. generated phenotypic data and contributed the samples for present study. All authors were involved in writing and reviewing the manuscript.

## 9. DATA AVAILABILITY STATEMENT

Researchers interested in using STR data must obtain approval from a Swedish Ethical Review Board and from the Steering Committee of the Swedish Twin Registry. Researchers using the data are required to follow the terms of an agreement containing a number of clauses designed to ensure protection of privacy and compliance with relevant laws. For further information, contact Patrik Magnusson.

## 11. SUPPORTING INFORMATION

Supporting Text Discussion, Materials and methods

Supporting Table 1. The HRG associated SNPs that are non-synonymous or located near to transcription start site.

Supporting Table 2. The results of stratified analysis by genotype of rs9898

Supporting Table 3. Enzymes used for digestion of human serum for MS/MS analysis

Supporting Table 4. Statistics for survival analyses from Figure 3 and Supporting Figure 7

Supporting Table 5. Summary of survival analyses

Supporting Figure 1. Study design

Supporting Figure 2. Difference between serum and plasma in sample sets 1 and 2

Supporting Figure 3. Protein profiles of HPA045005 in every sample set

Supporting Figure 4. Additional validation of molecular target

Supporting Figure 5. Manhattan plot and LocusZoom plot for BSI0137 antibody profile

Supporting Figure 6. Probabilistic identification of causal SNPs (PICS) analysis for the associated SNPs with HPA045005 and BSI0137

Supporting Figure 7. Survival curves for women and men, comparing two extreme quarters by HRG profiles

Supporting Figure 8. Detection of HRG peptides in human serum by MS/MS analysis

