## Supporting for "Profiles of circulating histidine-rich glycoprotein associate with chronological age and risk of all-cause mortality"

### Supporting Text

#### 1. Discussion

##### GDF15 and medication.

Other recent affinity proteomics approaches have reported age-associations of circulating proteins, for example, growth differentiation factor 15 (GDF15) as well as other proteins of the coagulation system (Tanaka et al., 2018). This study also acknowledged the need for further validation. It is noteworthy that circulating GDF15 can also be highly influenced by medications such as metformin (Gerstein et al., 2017) and the expression of the *GDF15* gene, also known as nonsteroidal anti-inflammatory drugs-activated gene (*NAG-1*), can be induced by other common drugs (Wang, Baek, & Eling, 2011).

#### 2. Samples and selections

##### a) Sample set 6 from IMPROVE cohort

IMPROVE is a multicenter, longitudinal, observational cohort study of individuals at high-risk of cardiovascular diseases, from seven recruiting centers in five European countries: Finland, Sweden, the Netherlands, France and Italy (Baldassarre et al., 2010). A total of 3,711 participants were enrolled between March 2004 and April 2005. Eligibility criteria included age from 55 to 79 years, presence of at least three cardiovascular risk factors and absence of symptoms of cardiovascular diseases. Cases denote individuals who suffered a vascular event from month 15 to month 36 during follow-up, while controls did not have any vascular event during the time. Forty-three cases and age-, sex- and center-matched controls were used in the present study.

##### b) Sample set 7 from SCARF study

The Stockholm Coronary Atherosclerosis Risk Factor (SCARF) study included survivors of a first myocardial infarction (MI) before the age of 60 years and age- and sex-matched control subjects free from the disease at enrollment, between January 1996 and December 2000 (Samnegård et al., 2005). Cases were patients admitted to the three hospitals in the northern part of Stockholm area with an ongoing cardiac event, and controls were recruited in parallel from the general population of the same residence area. The blood samples were collected 3 months after the index cardiac event.

##### c) Sample set 8 from CHAPS

In the Carlscrona Heart Attack Prognosis Study (CHAPS), 5292 human subjects were consecutively recruited and EDTA plasma samples were collected in a coronary intensive care unit during 1992 to 1996 (Odeberg et al., 2014). Of these 908 patients aged 30–74 years had at discharge had received the diagnosis of either myocardial infarction (527) or unstable angina (381). As control group, 948 patients aged 30–74 years were identified who were admitted with suspected acute coronary syndrome (ACS), but were subsequently diagnosed as non-ACS and, furthermore, were not diagnosed with stable CAD. From this group, forty-three ACS patients and an equal number of non-ACS controls were selected for a nested case-control discovery study. Those controls were analyzed in the present study. The activated partial thromboplastin time (aPTT) was analysed in Sodium Citrate blood samples using a routine diagnostic method on a Trombotrack instrument (Nycomed, Norway). The procedures for blood sampling and laboratory analyses followed the routines of the Department of Clinical Chemistry at Blekinge County Hospital and analyses were performed in the certified hospital laboratory using fresh samples collected at hospital admission. The hospital laboratory reference range with this assay was APTT 22-32 s.

**d) Sample set 9 from Karma study**

The Karolinska Mammography Project for Risk Prediction of Breast Cancer (KARMA; www.karmastudy.org) is a prospective population-based cohort (Gabrielson et al., 2017). Nearly 71,000 women were included in the cohort between 2011-2013. Women invited for mammography screening or clinical mammography at any of the four mammography units in Sweden were recruited into the cohort study. At study entry, information on a vast number of lifestyle factors was gathered through a web-based questionnaire. Plasma samples from non-diseased women were selected from mammographic breast density measurements (absolute volumetric density, $cm^3$), where 300 dense and 300 non-dense samples were matched on age (median 54, 40-73) and body mass index (median 24, 19-30).

**e) Sample selection for replication**

The subjects of the sets 2 to 9 comprised of 829 subjects from non-diseased control groups and 3,215 chosen to reflect Swedish population. The entire set of subjects were from 3 to 93 years old at blood draw. Blood samples had been prepared either as serum or plasma (Table 1, Supporting Figure 2). Hence, due to the plausible differences originating from these preparation types, twelve sera samples in the set 2 were excluded from the following meta analyses as the remaining samples in this set were plasma. The sample set 2 (216 subjects) was chosen from a population-based prospective cohort in Sweden (Almqvist et al., 2011) and the set 3 (2999 subjects) was from the twin cohort same as the set 1 (Lichtenstein et al., 2002; Magnusson et al., 2013), in which disease status was not considered during recruitment. Almost all (98.5%, all except forty-four) subjects in the set 3 were not overlapped with the set 1. Two other sample sets (set 4 and 5) included 100 subjects that were selected from cancer-related studies and derived from the same twin cohort as the set 1 (Lichtenstein et al., 2002; Magnusson et al., 2013). Besides one single subject, there was no overlap between these and the individuals analyzed during the discovery. Sample sets 6 to 9 (729 subjects) were from four independent studies (Table 1) (Baldassarre et al., 2010; Gabrielson et al., 2017; Odeberg et al., 2014; Samnegård et al., 2005).

#### 3. Suspension Bead Array Assays

##### a) Antibody selection

Antibodies from the Human Protein Atlas (HPA) with a concentration above 0.05 mg/ml and passing specificity assessment on planar protein microarrays (Sjoberg et al., 2012) were used for the analysis of sample sets 1 and 2. No other protein related criteria other than antibody availability at the time of the study were applied. For the analysis of sample sets 4, 5, 6, 7, and 8, antibodies were selected without considering target proteins. The antibodies for sample set 3 included antibodies that had been used and showed some potentially interesting results in other studies for various diseases. Two of them were HPA045005 from the HPA and BSI0137 (Batch ID 0137090310), which is a monoclonal mouse antibody from BioSystems International Kft (Guergova-Kuras et al., 2011). The antibodies for sample set 9 were chosen for a mammography study for its own interest (Bystrom et al., 2018).

##### b) Suspension Bead Array Procedure

Coupling of antibodies to magnetic beads (MagPlex, Luminex Corp) was performed as previously described (Byström et al., 2014) using 1.6 µg of a given antibody per 500 000 beads. For this purpose, antibodies were diluted using a liquid handler (EVO150, TECAN), beads were washed on a magnet using a plate washer (EL406, Biotek). Coupled beads were blocked and stored in buffer (Blocking reagent for ELISA, Roche) supplemented with ProClin (Sigma) at 4°C in the dark. Equal volumes of beads carrying different capture antibodies were mixed to create a suspension bead array (SBA). As described before, samples were labeled with biotin and heat-treated (Schwenk et al., 2010). In short, samples were centrifuged, diluted 1/10 in PBS using a liquid handler (SELMA, CyBio AG), labeled with 10 mg/ml NHS-biotin (Pierce), which was quenched using 0.5 M Tris-HCl. Labeled samples were diluted 1:50 in assay buffer (0.5% (w/v) polyvinyl alcohol and 0.8% (w/v) polyvinylpyrrolidone (Sigma) in 0.1% casein in PBS supplemented with 0.5 mg/ml rabbit IgG (Bethyl Laboratories) and heat-treated for 30 min at 56°C using thermocyclers. Then 45 µl of samples were added to 5 µl SBA, incubated overnight at room temperature with constant rotation at 650 rpm (Grant). Thereafter, beads were washed 3x in 100 µl PBS-T and 50 µl 0.4% PFA was added for 10 min. Beads were washed once again with PBS-T before streptavidin R-PE (Invitrogen) was diluted 1:750 in PBS-T and 50 µl was added for 20 min. Prior to analysis in FlexMap3D instrument (Luminex Corp), beads were washed 3x in 100 µl PBS-T. Median signal intensities (MFI) of each bead ID were used for subsequent data analysis.

##### c) Assay design

All 372 samples from sample sets 1 and 2 together were randomly allocated into wells in four 96-well plates. One sample from sample set 1 and one from sample set 2 were loaded into two more wells as a repeated control within a plate. Another sample in each cohort was transferred to two more wells of two different plates as a control to examine inter-plate variation. The data of each of those 4 samples was combined by taking mean of three measures. All the human materials were biotinylated together with four negative controls that contained only buffer. For the entire 19 assays for discovery stage, the samples were labeled two times.

The selected antibodies were divided into collections of 384 antibodies including positive and negative controls, anti-albumin and no antibody, respectively. These antibodies were then coupled onto beads and used to create a suspension bead array (SBA). For discovery, the selection of the affinity binders for one SBA was determined by technical reasons such as the available amount. Every antibody in an SBA was coupled with beads with a different color code as detailed together with the assay procedure in the Supporting Information and as described earlier (Byström et al., 2014). This assay provided protein profiles from up to 384 antibodies and 384 samples per batch.

**d) SBA data quality control and preprocessing**

Because an aliquot of mixed bead solution was suspended into each sample, all values of the samples that were seemingly failed within an assay (with 384 antibodies) were discarded rather than one measure of a sample for an antibody. These were samples 1) that had median bead counts lower than twenty, 2) that had median values of MFIs lower than the median of the negative controls (buffer only) in the same plate and assay, and 3) that were detected as an outlier by robust PCA using 'rrcov' R package (version 1.3-4)(Hubert, Rousseeuw, & Branden, 2005). The cutoff probability values in an outlier diagnostic plot were 0.025 for both score and orthogonal distance coordinates. Those deviating beyond the cutoffs in both coordinates were classified as outliers, setting alpha, the proportional tolerance, to 0.9.

The human samples were of two different types in terms of preparation method, plasma and serum. The two blood preparation types showed considerable dissimilarity, which was expected (Supporting Figure 2). Since such contrast was not of our research interest here, the data was split by the sample preparation type after quality control, when two types of samples were analyzed in the same assay plates.

The variation across sample plates was minimized by Multi-MA method, with the assumption that the mean of observed values for each antibody within a plate is same as those of the other plates (Hong, Lee, Nilsson, Pawitan, & Schwenk, 2016) . The means of log-transformed measures within each plate were positioned in a 4-dimensional space, in which each axis corresponded to one plate. The vector that goes through origin and (1, 1, 1, 1) is named A. The projection of each point onto the A axis is computed. All values for the point were shifted as much as the element vector on the corresponding plate axes of the projection. The proteomic data for pQTL analysis was pre-processed including technical controls and using probabilistic quotient normalization (PQN) (Kato et al., 2011) before the Multi-MA normalization.

**e) Data acquisition of replication sample sets**

Data of other replication samples (sample sets 3-9) were acquired using the same protocol with a few variations. For each original study for sample sets 4-8, the samples were distributed into plates together with patient samples. The 383 other antibodies selected for each of the intended studies were included in the assays. Experiment and data preprocessing were conducted together with those additional samples and antibodies. Data of disease-free controls and for HPA045005 were extracted from the processed full data sets.

#### 4. Sandwich immunoassays

To confirm the capture of HRG by HPA045005, a sandwich immunoassay was developed and used for the detection of a full-length recombinant HRG protein, which was a kind gift from Hanna Tegel and Johan Rockberg (AlbaNova University Center, KTH). A multiplexed SBA containing a library of HPA antibodies including HPA045005 and the anti-HRG antibody HPA054598 were used as capture reagents, as described previously (Häussler et al., 2019). An empty bead and one rabbit IgG bead were also included in the SBA for measuring background signal. For detection, the antibody HPA054598 was biotinylated as previously described (Dezfouli et al., 2014). The HRG protein was prepared in a four-fold serial dilution (500 ng/ml to 0.1 ng/ml) in PVXC buffer (0.1% casein, 0.5% (w/v) polyvinylalcohol, 0.8% (w/v) polyvinylpyrrolidone, Sigma- Aldrich) supplemented with 0.5 mg/ml purified rabbit IgG (Bethyl laboratories). The serial dilution was added in triplicate to a 96-well plate along with three control PVXC buffer wells. Five microliters of the SBA were incubated overnight with 45 µl protein standard. Following that, the beads were washed three times in PBS-T 0.05%. The biotinylated detection antibody was diluted to 1 µg/ml in PBS-T 0.05% and 25 µl was transferred to the beads (SELMA, Cybio). After 1.5 h incubation at room temperature, the beads were washed three times in PBS-T 0.05%. R-phycoerythring-labeled streptavidin (Invitrogen) was diluted 1:500 in PBS-T 0.05% and 50 µl incubated with the SBA for 20 min. Finally, beads were washed three times and measured in PBS-T with a FlexMap 3D instrument. Presence of detection antibody was confirmed with an anti-rabbit IgG bead.

#### 5. Protein and peptide microarray analysis

To determine the selectivity of the antibody binding, a set of 16,728 protein fragments from the Human Protein Atlas were used to generated microarrays with 21,120 features corresponding representing 12,412 unique Ensemble Gene IDs according to a previous protocol (Sjoberg et al., 2012). The array contained > 12,400 protein fragments of a length between approximately 20-150 residues and included the antigen used to generate HPA045005

We also aimed at determining the binding of HPA045005 to peptides representing its antigen using high density peptide arrays from NimbleGen using a previously described protocol. (Forsstrom et al., 2014). The array a set of 12-mer peptides with 11 residue overlap representing the antigen did not reveal a significantly prominent recognition of these peptides above background.

#### 6. Mass spectrometry analysis

An LC-MS/MS proteomics experiment was carried out to verify the findings from the affinity proteomics experiments orthogonally. A pool serum samples from 3 males and 2 females was diluted 5 times with 1x PBS. Sodium deoxycholate (SDC) was added to a final concentration of 1% (w/v) and proteins reduced and alkylated for 10 min at 96°C in 10 mM dithiothreitol and for 30 min at room temperature in the dark in 50 mM chloroacetamide for 30 min. The sample was diluted with 1x PBS to the final SDC concentration of 0.1% (w/v) and split into 5 tubes so each tube contained 1 uL of raw plasma. Protein digestion was performed using 5 different enzymes (0) to increase the peptide coverage by data dependent acquisition (Tsiatsiani & Heck, 2015). Formic acid was added to quench the enzymatic reaction to the final concentration of 1% (v/v) and SDC was let to precipitate for 30 min at room temperature. Samples were centrifuged at 16 400RCF for 5 min and subjected to LC-MS/MS analysis.

Samples were analyzed using the UltiMate 3000 capillary-liquid chromatography system (Thermo Scientific) with an EASY-Spray ion source connected to Q Exactive HF (Thermo Scientific) mass spectrometer. In total, 3.5 µg of peptides were loaded onto PepMap100 trap column (300 µm × 5 mm, C18, 5µm, 100 Å, Thermo Scientific), washed 5 min at 15 µl/min with 100% of Solvent A (3% ACN, 97% $H_20$, 0.1% FA) and separated with PepMap RSLC C18 (150 µm x 15 cm, 2 µm, 100 Å, Thermo Scientific) analytical column using linear 60 min gradient of 1-32% Solvent B (95% ACN, 5% $H_20$, 0.1% FA) at a flow rate of 3.6 µl/min. The analytical and trap columns were kept at 55°C by the in-source temperature controller and 40°C by the column oven temperature controller respectively. The MS analysis was performed using a Top10 method starting with an MS1 scan performed at resolution of 60,000 (mass range 350–1,200 m/z, AGC $3e^6$, max IT 100 ms) and followed by ten consecutive MS2 scans at resolution of 30,000 (AGC $2e^5$, max IT 105 ms) with normalized collision energy set to 28.

Resulting raw files were searched using MaxQuant version 1.6.1.0 (Cox & Mann, 2008) with the built-in search engine Andromeda against wild type sequence of HRG and HRG sequence that contained the four SNP variants (rs10770, rs9898, rs2228243, rs1042464). Search parameters were set to peptide length ranging 7-25 amino acids, enzyme specificity was according to the used enzyme for digestion (0) with maximum of two mis-cleavages, 4.5 ppm match tolerance for precursor ions and 20 ppm for fragment ions with 1% false discovery rate on both the peptide and protein level. Carbamidomethylation on cysteine was set as a fixed modification.

The assay was able to detect one region that included the N493I dbSNP (rs1042464). Additionally, data from the Peptide Atlas was used to look for evidence of the missing SNPs. It confirmed both that the rs1042464 variant has been observed and yield useful proteotypic peptides suitable for LC-MS/MS analysis as well as the three other SNPs are never found (rs10770, rs2228243) or very unlikely to be observed (rs9898) (Supporting Figure 8). This is probably due to the properties of peptides being unsuitable for LC-MS/MS analysis in terms of bad ionization or due to the matrix effects of plasma proteome.

#### Supporting Figures

**Supporting Figure 1.**

Study design. This illustration describes the steps of the present investigation.

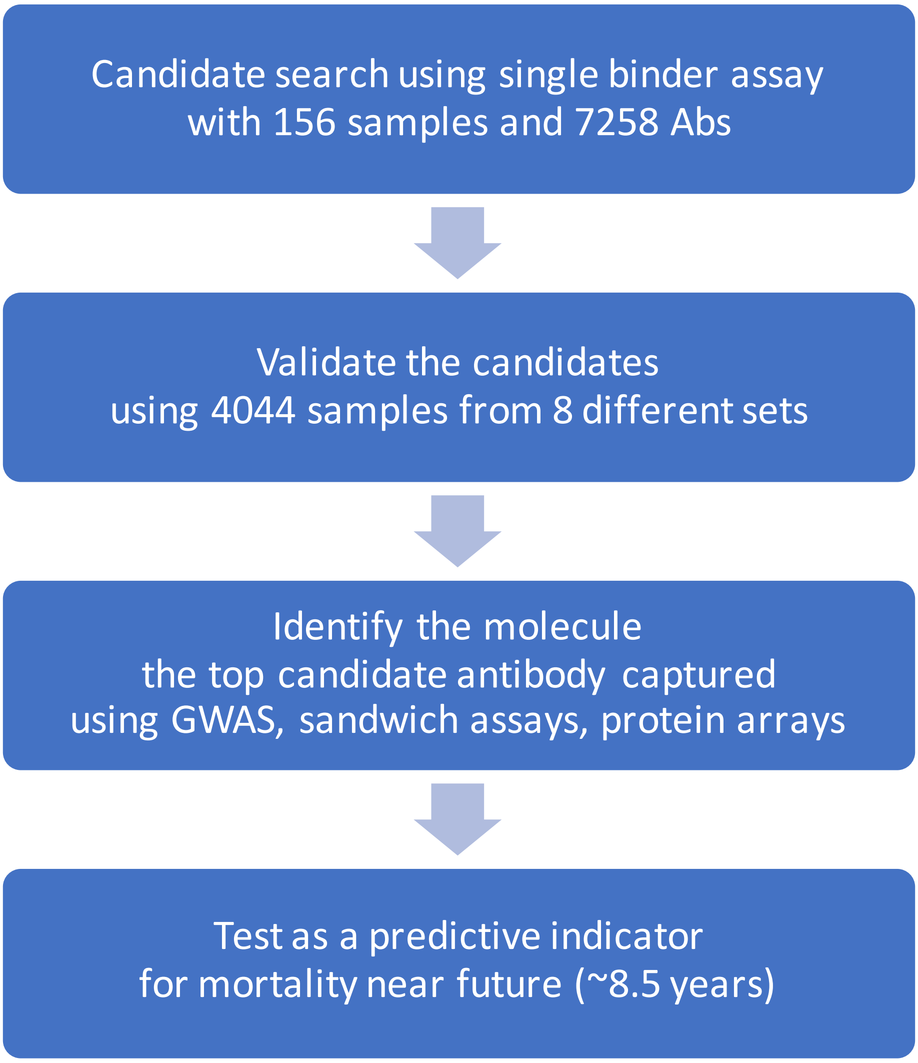

**Supporting Figure 2.**

Difference between serum and plasma in sample sets 1 and 2. Each plot displays the first 2 principal components of the MFIs of one assay, where the profiles of a set of 384 antibodies were obtained. The points of the scatter plot were colored by sample preparation types, such as plasma and serum.

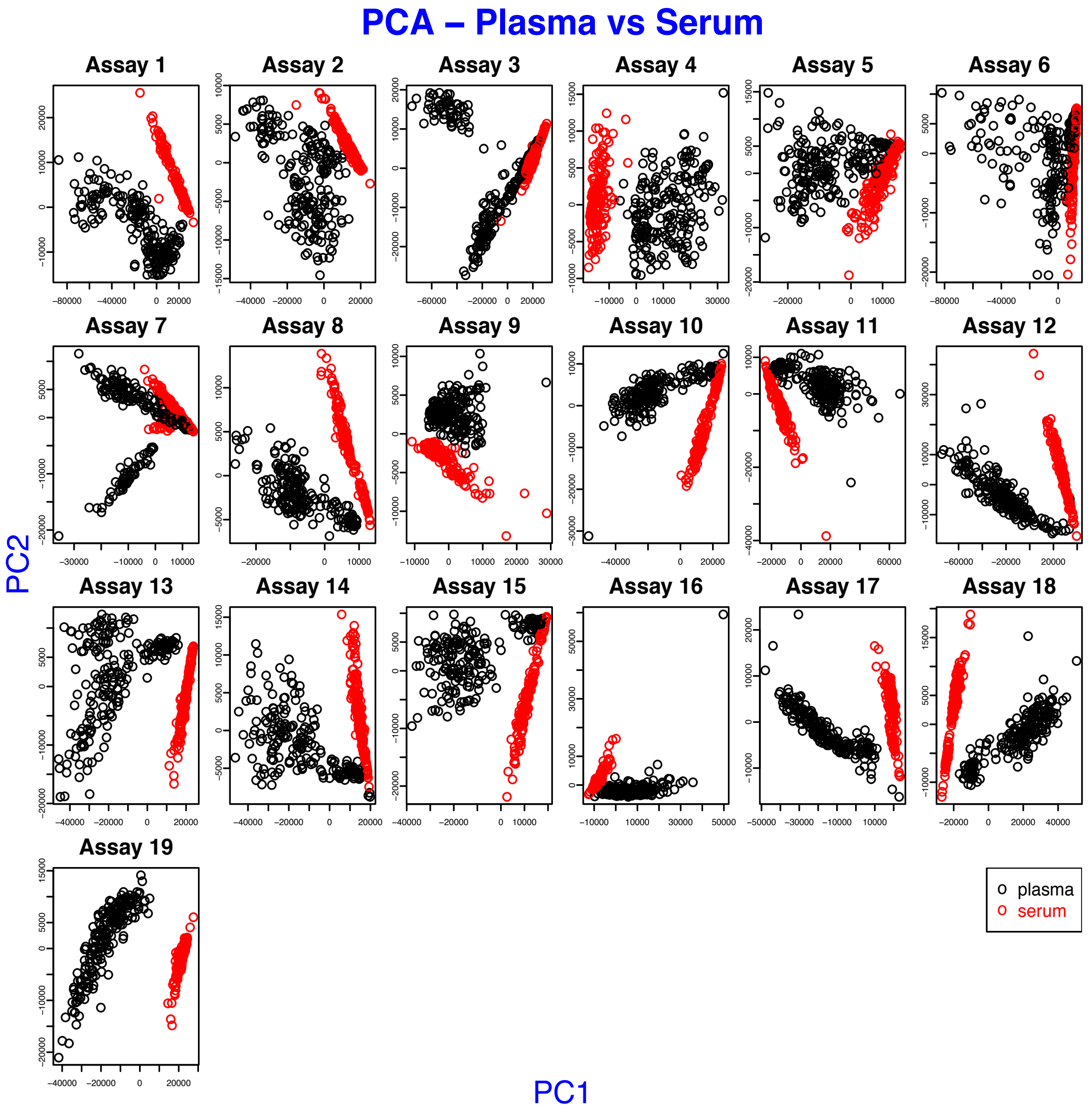

**Supporting Figure 3.**

Protein profiles of HPA045005 in every sample set. The solid lines illustrate the trends estimated by linear regression, and the shades around them show 95% confidence intervals of fitted values. At the bottom right corner, all trend lines were overlapped to present the increasing trends in all 9 sample sets.

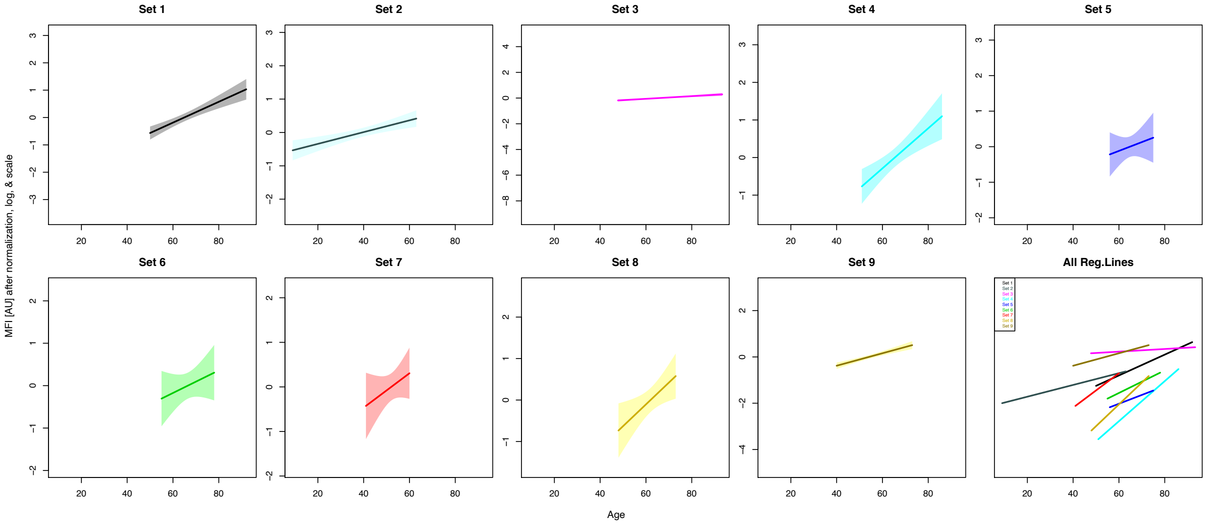

**Supporting Figure 4.**

Additional validation of molecular target (A) Comparative sandwich immunoassays for HRG. To confirm the molecular binding of HPA045005 to HRG, dilution of spiked-in HRG protein levels were studied. Different colors represent additional capture antibodies, such as the anti-HRG binder HPA054598. All were used for a multiplexed assessment of HRG binding. For the detection a labelled version of HPA054598 was used. (B) High density protein array analysis of HPA045005. The signal intensities from the interaction of the antibody with any of the 16,728 immobilized unique protein fragments on the glass slide are illustrated. For the antibody HPA045005, one peak was exclusively observed for the protein fragment that had been employed for the production of the antibody (red bar), as well as for the positive controls detecting the presence of the rabbit IgG antibody (orange and green bars).

A.

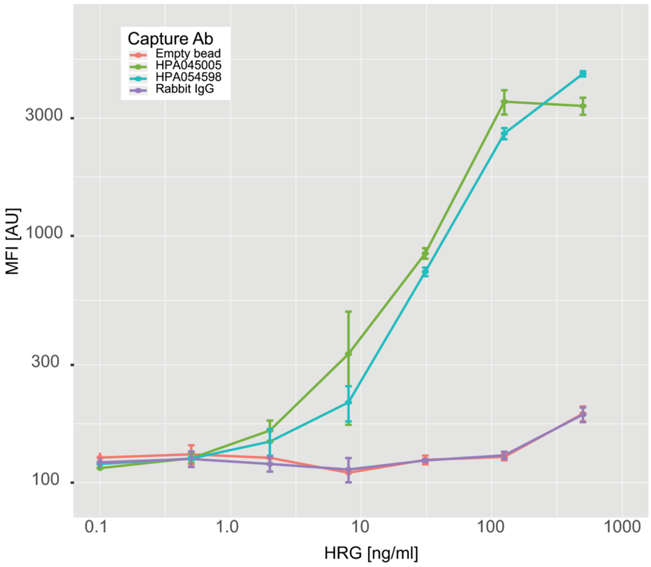

B.

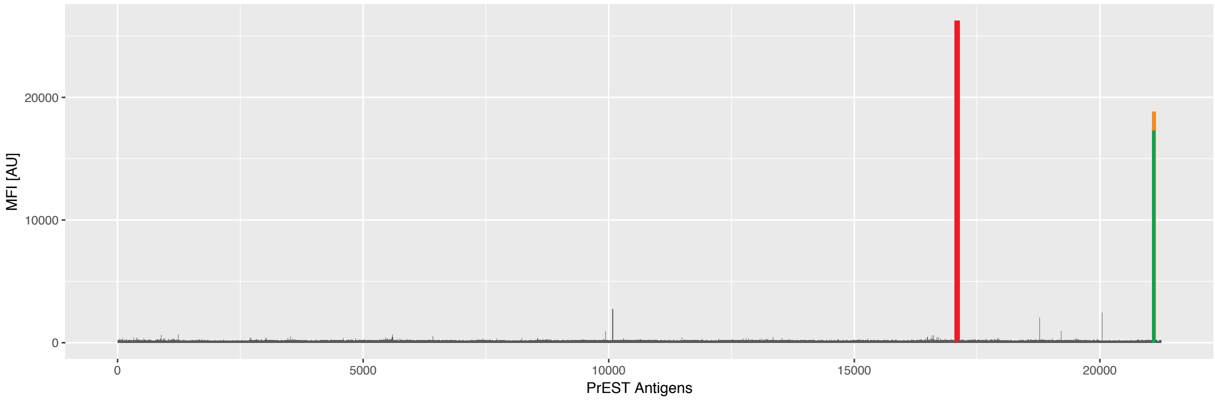

**Supporting Figure 5.**

Manhattan plot and LocusZoom plot for BSI0137 antibody profile. Because the P-values of the top SNPs in the GWAS for BSI0137 was too low, the computation tool for this analysis, PLINK, gave us 0s. The values were manually set to $1 \times 10^{-320}$ in (A) or $1 \times 10^{-350}$ in (B) for this visualization.

A.

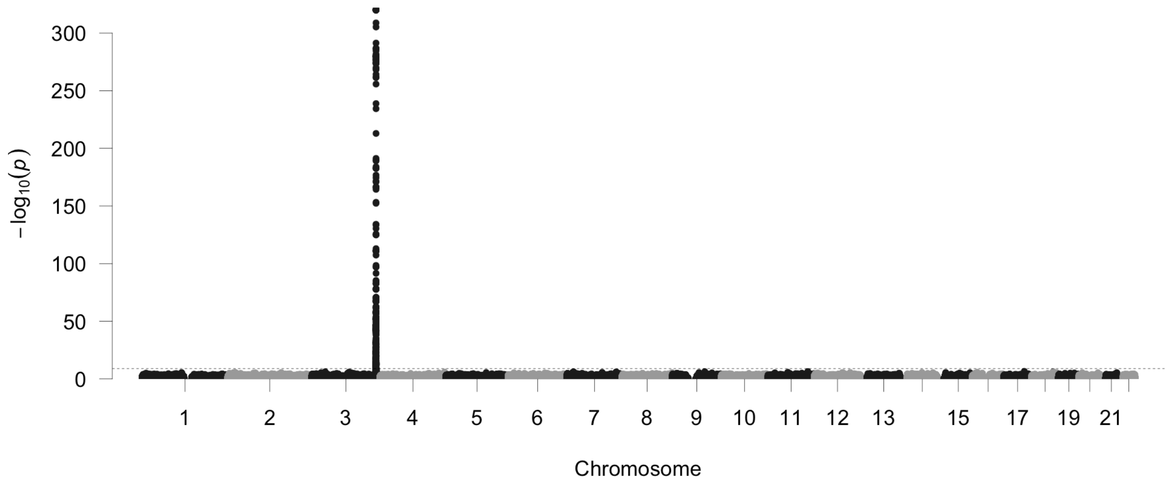

B.

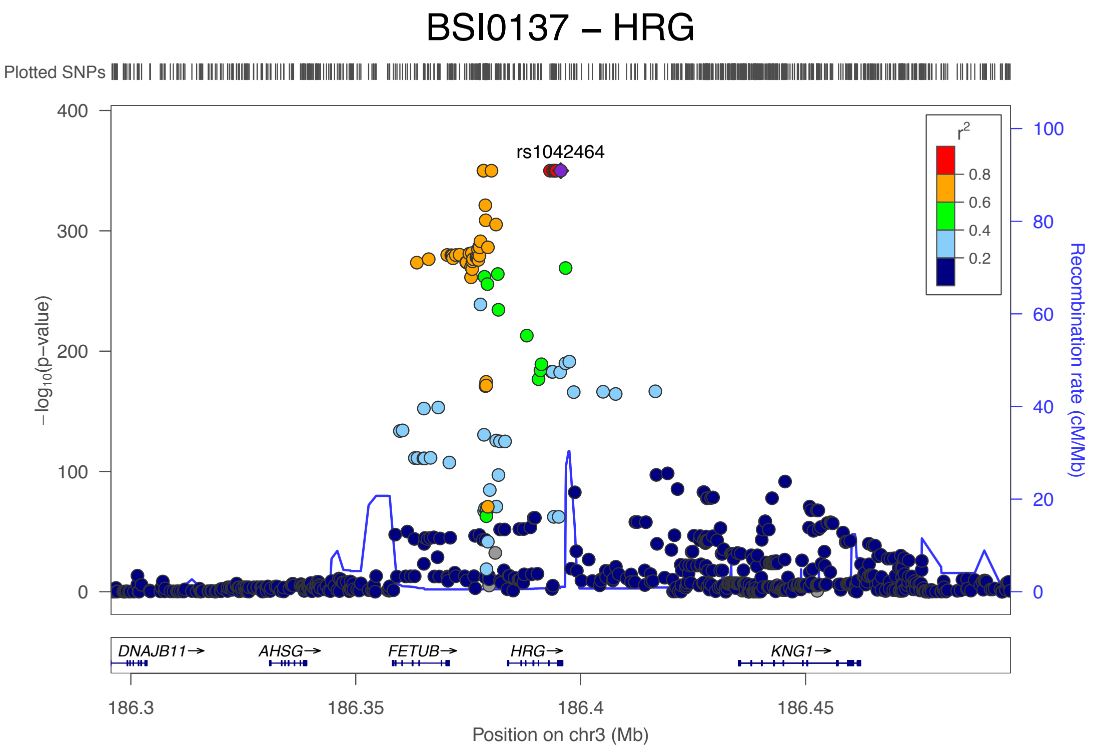

**Supporting Figure 6.**

Probabilistic identification of causal SNPs (PICS) analysis for the associated SNPs with HPA045005 and BSI0137. The top two figures display the distribution of -log of P-values obtained from the linear regression model for the effects of the genotype of rs1042464 in 100000 permutations, assuming rs9898 was causal for individual antibody profiles. The bottom figures show the results of rs9898 assuming rs1042464 was causal genetic variant. The green vertical lines indicate the observed significances.

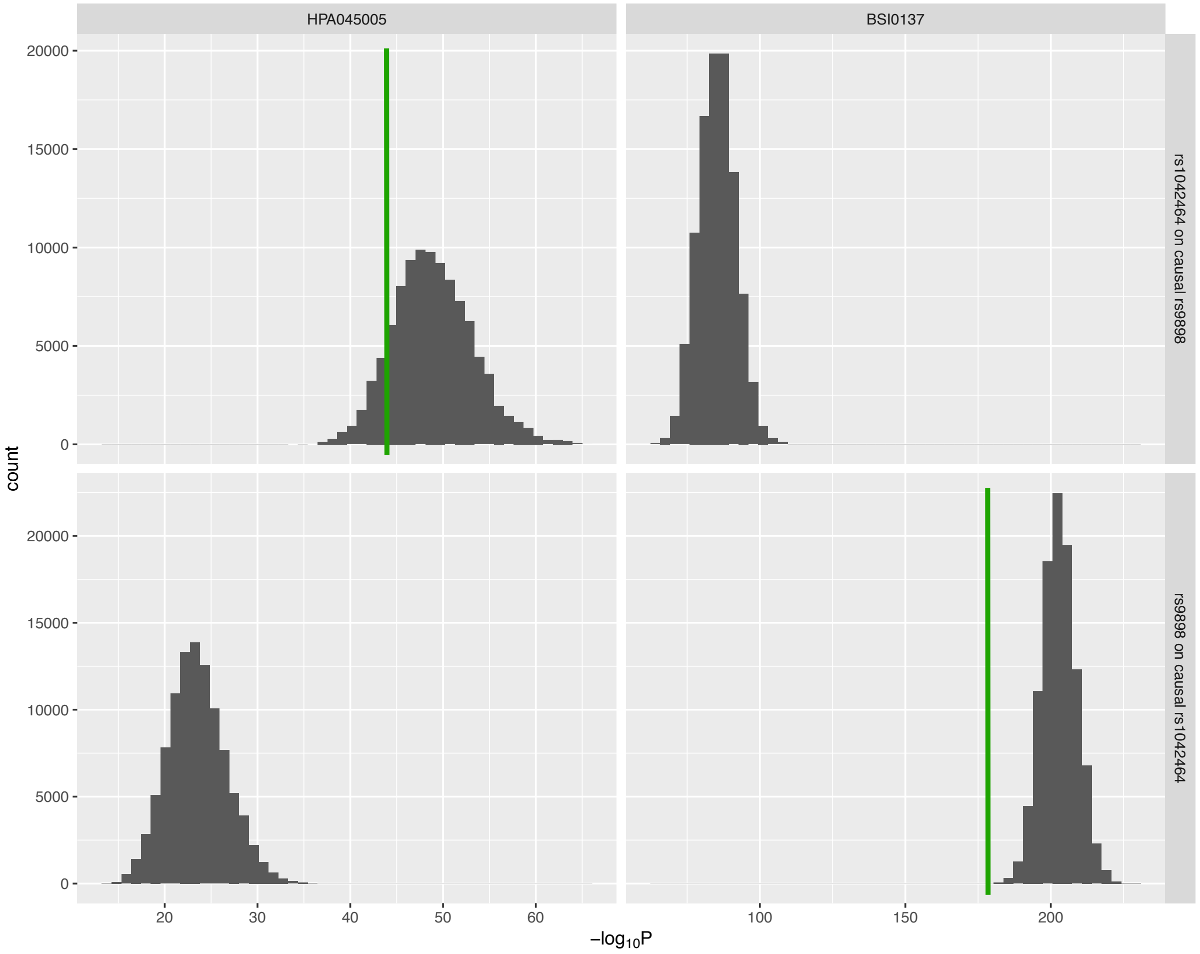

**Supporting Figure 7.**

Survival curves for women and men, comparing two extreme quarters by HRG profiles.

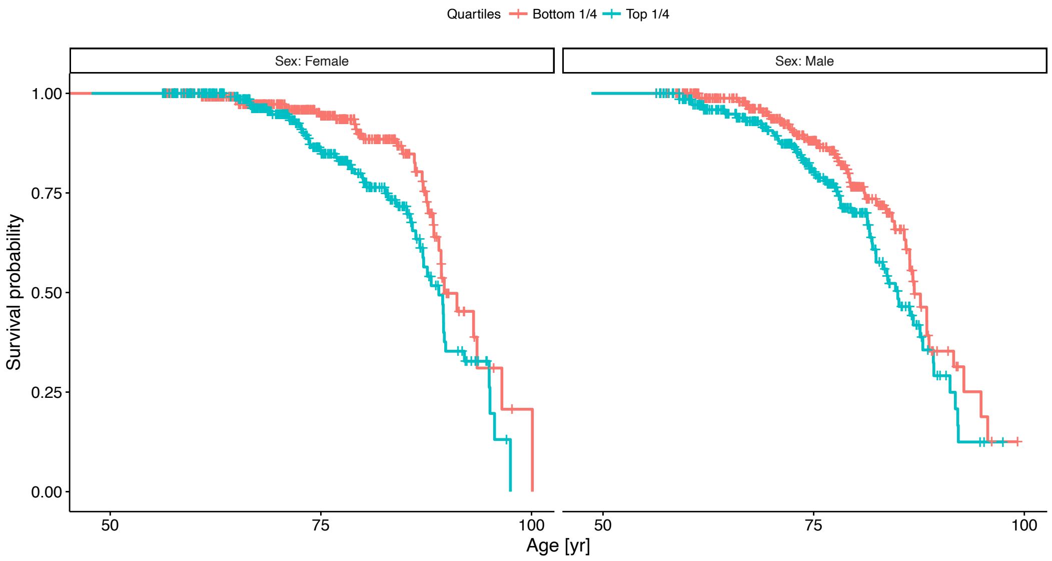

**Supporting Figure 8.**

Peptides from HRG detected in serum samples using a bottom-up proteomic experimental workflow and liquid chromatography tandem mass spectrometry for read-out. Top panel: Five different enzymes (y-axis; GluC, LysN, LysC, Chymotrypsin and Trypsin) and the peptide amino-acid sequence coverage (x-axis) of their experimental peptides. Four SNP variants (rs10770, rs9898, rs2228243 and rs1042464) have been indicated by vertical black lines. Bottom panel: Experimental peptides reported by the Peptide Atlas (accessed 2017-05-16). The peptide amino acid sequence coverage (x-axis) is visualized in relation to experimental peptides for each experiment (y-axis). Highlighted in this example, rs9898 has only a few theoretical peptide sequences that can be observed with the bottom up method (due to sequence length) using this set of enzymes. Either one tryptic peptide with one missed cleavage NCPRHHFPR (identified once highlighted in the figure) or the chymotryptic peptide SCRNCPRH (based on low enzymatic specificity). For both panels, blue indicates the most abundant protein form, and red is for any variation from the form.

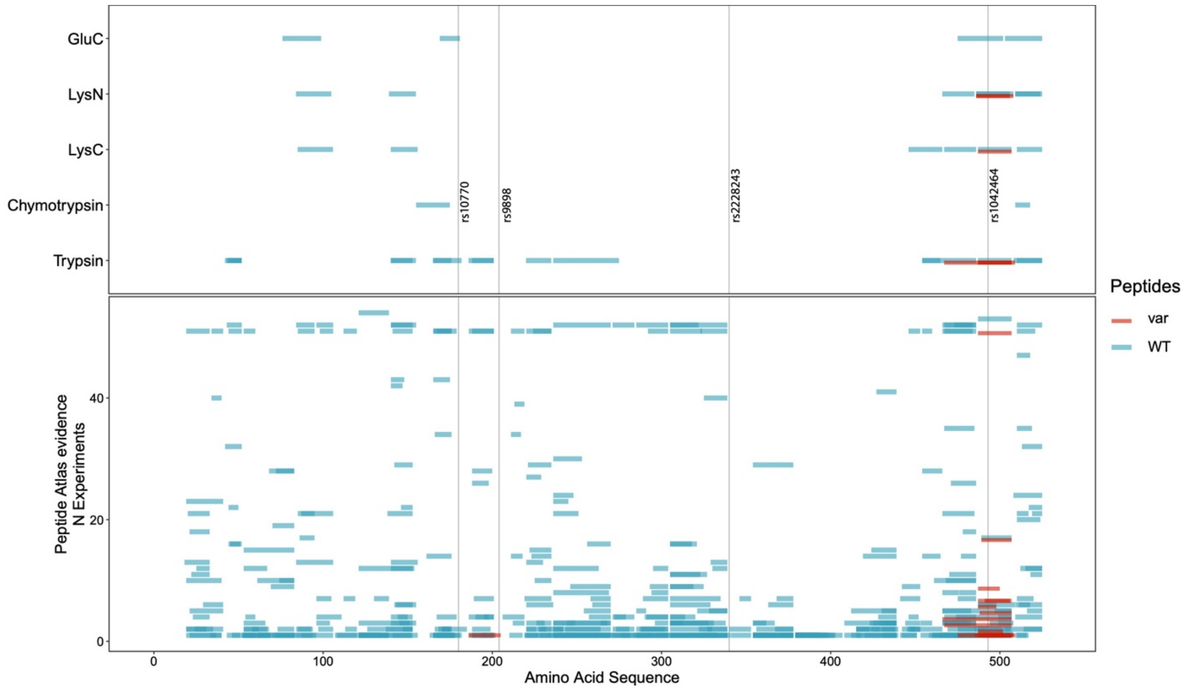

### Supporting Tables

##### Supporting Table 1.

The HRG associated SNPs that are non-synonymous or located near to transcription start site.

| SNP | P value† | MAF‡ | Coefficient (β) † | Amino-acid variants [Codon] | Position in precursor |
|---|---|---|---|---|---|
| rs9898 | $2.4 \times 10^{-97}$ | 0.32 | 0.15 | Pro : Ser [**C**CC:**T**CC] | 204 |
| rs1042464 | $1.9 \times 10^{-44}$ | 0.47 | 0.10 | Asn : Ile [A**A**T:A**T**T] | 493 |
| rs2228243 | $1.6 \times 10^{-34}$ | 0.20 | 0.11 | His : Arg [C**A**T:C**G**T] | 340 |
| rs10770 | $3.9 \times 10^{-26}$ | 0.10 | 0.12 | Ile : Thr [A**T**C:A**C**C] | 180 |
| rs3890864 | $6.6 \times 10^{-46}$ | 0.16 | 0.13 | close to TSS‡ | |
| rs56376528 | $8.3 \times 10^{-46}$ | 0.16 | 0.13 | | |

† Results from the linear regression for allele frequency and HRG level

‡ MAF = minor allele frequency, TSS = transcription start site

##### Supporting Table 2.

The results of stratified analysis of HPA045005 by genotype of rs9898.

| Genotype | Amino acid | N | HR* (hazard ratio) | HR P value* | Coefficient (β) † | LM P value† |
|---|---|---|---|---|---|---|
| CC | Proline | 1094 | 1.32 | 0.012 | 0.0007 | 0.23 |
| CT | Pro, Ser | 970 | 1.30 | 0.015 | 0.0027 | $6.9 \times 10^{-5}$ |
| TT | Serine | 243 | 1.19 | 0.27 | 0.0050 | $1.6 \times 10^{-3}$ |

* Cox proportional hazards models using age-adjusted HRG level from HPA045005.

† Linear regression models for age and HRG level from HPA045005. The coefficients are the estimates of how much log of the HRG levels increase per year of age.

**Supporting Table 3.**

Enzymes used for digestion of plasma pool for MS/MS analysis

| Enzyme | Enzyme : Substrate ratio | Digestion time (hours) |
|---|---|---|
| **GluC** | 1:50 | 16 |
| **LysN** | 1:100 | 16 |
| **LysC** | 1:100 | 16 |
| **Chymotrypsin** | 1:50 | 16 |
| **Trypsin** | 1:50 | 1 |

**Supporting Table 4.**

Statistics for the survival analyses presented in Figure 3 and Supporting Figure 7. Ages were grouped into 5-year intervals. The number of individuals who entered each age-interval and died or censored within the interval was clarified.

| Age-interval | Female | | | | | | | |
|---|---|---|---|---|---|---|---|---|
| | Top 1/4 | | | | Bottom 1/4 | | | |
| | $N_{ft}^*$ | $D_{ft}^*$ | $C_{ft}^*$ | $LE_{ft}^†$ | $N_{fb}^*$ | $D_{fb}^*$ | $C_{fb}^*$ | $LE_{fb}^†$ |
| **(45,50]** | 13 | 0 | 0 | 87.4 | 13 | 0 | 0 | 91.0 |
| **(50,55]** | 53 | 0 | 0 | 87.4 | 54 | 0 | 0 | 91.0 |
| **(55,60]** | 117 | 0 | 28 | 87.5 | 120 | 0 | 31 | 91.0 |
| **(60,65]** | 176 | 1 | 36 | 87.7 | 156 | 2 | 46 | 91.1 |
| **(65,70]** | 218 | 6 | 82 | 88.1 | 211 | 1 | 75 | 91.2 |
| **(70,75]** | 192 | 12 | 73 | 88.8 | 194 | 4 | 60 | 91.4 |
| **(75,80]** | 135 | 7 | 56 | 89.9 | 158 | 4 | 85 | 91.9 |
| **(80,85]** | 91 | 5 | 49 | 91.5 | 88 | 3 | 42 | 92.8 |
| **(85,90]** | 45 | 16 | 14 | 93.7 | 47 | 12 | 23 | 94.3 |
| **(90,95]** | 16 | 2 | 10 | 96.5 | 13 | 3 | 6 | 96.6 |
| **(95,100]** | 4 | 3 | 1 | 100 | 4 | 1 | 2 | 99.8 |
| **(100,105]** | 0 | 0 | 0 | | 1 | 1 | 0 | |

| Age-interval | Male | | | | | | | |
|---|---|---|---|---|---|---|---|---|
| | Top 1/4 | | | | Bottom 1/4 | | | |
| | $N_{mt}^*$ | $D_{mt}^*$ | $C_{mt}^*$ | $LE_{mt}^†$ | $N_{mb}^*$ | $D_{mb}^*$ | $C_{mb}^*$ | $LE_{mb}^†$ |
| **(45,50]** | 8 | 0 | 0 | 83.9 | 4 | 0 | 0 | 86.6 |
| **(50,55]** | 43 | 0 | 0 | 84.1 | 43 | 0 | 0 | 86.7 |
| **(55,60]** | 89 | 1 | 18 | 84.3 | 87 | 0 | 13 | 86.8 |
| **(60,65]** | 131 | 3 | 39 | 84.7 | 138 | 1 | 44 | 87.0 |
| **(65,70]** | 190 | 5 | 56 | 85.4 | 178 | 6 | 58 | 87.4 |
| **(70,75]** | 177 | 15 | 55 | 86.4 | 179 | 8 | 49 | 88.1 |
| **(75,80]** | 136 | 11 | 72 | 88.0 | 148 | 12 | 74 | 89.2 |
| **(80,85]** | 63 | 13 | 26 | 90.0 | 72 | 6 | 37 | 90.9 |
| **(85,90]** | 28 | 6 | 15 | 92.7 | 34 | 10 | 15 | 93.2 |
| **(90,95]** | 8 | 4 | 2 | 96.0 | 10 | 3 | 4 | 96.1 |
| **(95,100]** | 2 | 0 | 2 | 99.8 | 3 | 1 | 2 | 99.8 |
| **(100,105]** | 0 | 0 | 0 | | 0 | 0 | 0 | |

* $N_x$ = the number of individuals entering the age-interval, $D_x$ = the number of dead subjects within the age interval, and $C_x$ = the number of individuals censored within the age interval.

† $LE_x$ = Life expectancy of the individual who is entering the age interval, i.e. lower bound age (e.g. 70 for (70,75] interval), assuming time-to-event follows Weibull distribution.

**Supporting Table 5.**
Summary of survival analyses

| Predictor* | Samples | N | Nr of deaths | Covariates | Hazard ratio (95% CI) | P value |
|---|---|---|---|---|---|---|
| **HRG** | All | 2973 | 362 | Sex | | $1.13 \times 10^{-4}$ |
| **BSI0137** | All | 2973 | 362 | Sex | | 0.57 |
| **std_HRG**[§] | All | 2973 | 362 | Sex | 1.25 (1.12-1.39) | $7.41 \times 10^{-5}$ |
| **std_HRG**[§] | All | 2307 | 295 | Sex, rs9898 | 1.31 (1.14-1.49) | $9.02 \times 10^{-5}$ |
| **std_HRG**[§] | Females | 1602 | 160 | | 1.36 (1.16-1.60) | $2.13 \times 10^{-4}$ |
| **std_HRG**[§] | Males | 1371 | 202 | | 1.15 (0.99-1.33) | 0.059 |
| **1st vs. 4th quartiles** | 2 quartiles | 1488 | 188 | Sex | 1.53 (1.29-2.30) | $4.60 \times 10^{-3}$ |
| **CRP** | All | 2971 | 362 | Sex | 1.07 (1.01-1.14) | 0.023 |
| **age-adjusted CRP** | All | 2971 | 362 | Sex | 1.01 (1.00-1.02) | 0.023 |
| **std_HRG**[§] | All | 2971 | 362 | CRP, Sex | 1.24 (1.11-1.38) | $1.20 \times 10^{-4}$ |
| **std_HRG**[§] | All | 2973 | 362 | Glucose, Sex | 1.25 (1.12-1.40) | $4.40 \times 10^{-5}$ |
| **std_HRG**[§] | All | 2970 | 361 | HbA1c, Sex | 1.25 (1.13-1.40) | $4.05 \times 10^{-5}$ |

[*] HRG profiles obtained by HPA045005.

[§] standardized HRG values by linear regression and scaling.
